# Non-canonical autophagy in dendritic cells restricts cross-presentation and anti-tumor immunity

**DOI:** 10.1101/789867

**Authors:** Payel Sil, Fei Zhao, Ginger W. Muse, Sing-Wai Wong, Joseph P. Kolb, Laura Miller DeGraff, Charles J. Tucker, Erica Scappini, Annelise G. Snyder, Sara Grimm, Andrew Oberst, Jennifer Martinez

**Affiliations:** Immunity, Inflammation, and Disease Laboratory, National Institute of Environmental Health Sciences (NIEHS), National Institutes of Health (NIH), Research Triangle Park, NC 27709, USA; Signal Transduction Laboratory, Fluorescence Microscopy, and Imaging Center, National Institute of Environmental Health Sciences (NIEHS), National Institutes of Health (NIH), Research Triangle Park, NC 27709, USA; Department of Immunology, University of Washington, Seattle, WA 98109, USA; Division of Intramural Research, National Institute of Environmental Health Sciences (NIEHS), National Institutes of Health (NIH), Research Triangle Park, NC 27709, USA

**Keywords:** autophagy, Rubicon, LC3-associated phagocytosis, cross-presentation, inflammation

## Abstract

Major Histocompatibility Complex I (MHC-I) molecules classically present peptides derived from endogenous antigens, but exogenous antigens can also gain access to the MHC-I machinery in dendritic cells (DCs), which can activate antigen-specific CD8^+^ T cells. This process, termed cross-presentation, can be triggered by the uptake of dying autologous cells, including tumor cells, by DCs. The molecular mechanisms that underlie efficient cross-presentation remain largely uncharacterized, and an improved understanding of these mechanisms might reveal novel strategies for anti-tumor therapies. Rubicon (RUBCN) is a molecule required for LC3-associated phagocytosis (LAP), but dispensable for canonical autophagy, and mice lacking this protein develop an autoimmune inflammatory pathology with age. Here, we demonstrate that *Rubcn*-deficient DCs have increased retention of engulfed cellular cargo in immature phagosomes resulting in increased phagosome-to-cytosol escape and antigen access to proteasome-mediated degradation. As a result, mice selectively lacking *Rubcn* in DCs mount stronger tumor antigen-specific CD8^+^ T cell responses and exhibit decreased tumor burden compared to wild type littermates. These findings identify LAP as a key regulator of cross-presentation and suggest that targeting RUBCN might represent a novel strategy for anti-tumor therapy.

## Introduction

The ability of the immune system to distinguish between self and non-self is integral to maintaining homeostasis and eradicating exogenous threats. Antigen-presenting cells (APCs), specifically DCs, educate the host’s adaptive immune response in a controlled and antigen-specific manner. CD8^+^ T cells are activated by peptides generated from intracellular antigens and presented in the context of MHC-I. Peptides derived from healthy self-antigens presented to CD8^+^ T cells in the periphery elicit a minimal or absent response, as CD8^+^ T cells expressing T cell receptors (TCRs) that recognize self-antigens undergo negative selection in the thymus prior to their population of the periphery (Rock et al., 2016). However, when cells are infected by virus or are transformed by an oncogene, they can produce proteins that are not considered self, such as virus-encoded proteins, and the presentation of these aberrant antigens via MHC-I activates a robust CD8^+^ T cell response (Hansen and Bouvier, 2009). Therefore, the MHC-I pathway represents an evolutionarily conserved mechanism that serves as a quality control mechanism for endogenous proteins and viral defense (Kaufman, 2018).

As the activation of CD8^+^ T cells is largely limited to their localization within secondary lymphoid tissues, they must rely on the migration of patrolling APCs to deliver antigens to them (Hansen and Bouvier, 2009). Classically, exogenous antigens are sensed as foreign, phagocytosed, processed by the endosomal-lysosomal network, and loaded onto MHC-II molecules to stimulate antigen-specific activation of CD4^+^ T cells (Rock et al., 2016). Cross-presentation is a mechanism by which DCs can traffic exogenous antigens, such as autologous dying cells, acquired during surveillance to the MHC-I pathway for the activation of the CD8^+^ T cell response (Cruz et al., 2017). Specific subsets of DCs, such as CD8α^+^ DCs, are the primary executors of cross-presentation (Pooley et al., 2001), as they retain phagocytosed material in less degradative compartments for longer periods than other phagocytes, allowing for limited proteolysis and prolonged preservation of antigen integrity (Belizaire and Unanue, 2009). Macrophages, for example, quickly degrade intraphagosomal cargo via efficient maturation of the endocytic compartments, therefore limiting their cross-presentation capacity (Savina and Amigorena, 2007). Cross-presentation is specifically critical for the immunologically mediated clearance of tumors, and mice deficient in cross-presentation fail to generate a tumor-specific CD8^+^ T cell response and display significantly increased tumor burden (Cruz et al., 2017; Fehres et al., 2014; Huang et al., 1994; Wolfers et al., 2001). Thus, the efficacy of MHC-I to mediate delivery of both self-made and exogenously acquired antigens to CD8^+^ T cells highlights the importance of cytotoxic T cell activity in mitigating a multitude of pathological threats.

Exogenous antigens can access the MHC-I machinery via two pathways – the cytosolic pathway or the vacuolar pathway. In the cytosolic pathway, internalized cargo can escape the phagosome and enters the cytosol, wherein it is subjected to proteasome-mediated degradation into peptides. These peptides can then be transported to the ER by transporter associated with antigen processing (TAP1/2), where further processing by ER aminopeptidase 1 (ERAP1) and eventual loading onto the MHC-I molecule occurs. Peptide-MHC-I complexes are then transported to the Golgi apparatus and shuttled to the plasma membrane. In the vacuolar pathway, however, peptide generation, processing, and loading onto MHC-I molecules occurs exclusively within endocytic compartments in a proteasome- and TAP1-independent manner (Cruz et al., 2017; Fehres et al., 2014; Joffre et al., 2012).

Recent work has highlighted the activity and phagosomal trafficking of Sec22b protein during both the cytosolic and vacuolar pathways of cross-presentation, and the absence of Sec22b results in compromised cross-presentation (Alloatti et al., 2017; Cebrian et al., 2011). In DCs, Sec22b translocates to the cargo-containing phagosome and regulates the transport of ER and ER-Golgi intermediate compartment (ERGIC) proteins to the phagosome. Mechanistically, Sec22b-mediated recruitment of ERGIC proteins promotes the export of intraphagosomal cargo to the cytosol and delays maturation of the cargo-containing phagosome (Alloatti et al., 2017; Cebrian et al., 2011). In some cases, proteasome-generated peptides can be re-introduced to the phagosome via Sec22b-mediated delivery of ERGIC proteins, like TAP1/2, and signaling via Toll-like receptors (TLRs) during cross-presentation can facilitate the translocation of MHC-I to the phagosome, where the peptide-MHC-I complexes are assembled and transported to the plasma membrane independent of the Golgi apparatus (Nair-Gupta et al., 2014).

Despite recent advances in our understanding of cross-presentation, neither the relative contributions of each pathway nor the signals that mediate the recruitment and activity of key components are fully understood. We and others have described a unique form of non-canonical autophagy that couples phagocytosis to the autophagy machinery, termed LAP (Martinez et al., 2011a; Sanjuan et al., 2009). LAP, but not canonical autophagy, requires the activity of Rubicon (RUBCN), a protein that binds and regulates both the Class III PI3K complex and the NOX2 complex, both of which are essential for LAP (Martinez et al., 2015). Cargo-containing phagosomes are decorated with lipidated LC3 to form the LAPosome, which can then fuse to lysosomes for proper processing and degradation of the phagocytosed material (Martinez et al., 2011a). *In vitro* studies reveal that *Rubcn^-/-^*macrophages phagocytose normally, but fail to translocate LC3-II to the phagosome. This defect results in cargo persisting in immature phagosomal compartments and increased production of pro-inflammatory mediators, such as IL-6 and lL-12 (Martinez et al., 2011a; Martinez et al., 2016b). As one of the most ubiquitous sources for cross-presented antigens is dying cells and engulfment of dying cells during efferocytosis is a trigger for LAP, we hypothesized that LAP played a role in regulating cross-presentation.

*Rubcn^-/-^* mice are a valuable tool to allow the study of LAP-dependent mechanisms and pathologies without confounding deficits in the canonical autophagy pathway (Martinez et al., 2016b; Martinez et al., 2015). Broadly, LAP is a regulator of immunotolerance during the clearance of dying cells (efferocytosis), and *Rubcn^-/-^*mice develop systemic lupus erythematosus (SLE)-like autoimmunity and autoinflammation with age, which is associated with defective processing of dying cells (Martinez et al., 2016b). Interestingly, aged *Rubcn^-/-^*mice exhibit increased levels of activated CD8^+^ T cells in their periphery, suggesting that LAP deficiency could confer an increased capacity for MHC-I cross-presentation (Cunha et al., 2018; Martinez et al., 2016b). Here, we show, both *in vivo* and *in vitro*, that *Rubcn^-/-^*DCs can cross-present antigens more efficiently than wild type DCs, resulting in increased CD8^+^ T cell proliferation and activation. We also demonstrate that the engulfed cargo persists within less degradative phagosomes in the absence of RUBCN. These phagosomes are enriched for RAB5A and Sec22b, and phagosome-to-cytosol escape is significantly increased in *Rubcn^-/-^* DCs, compared to wild type DCs. This increased capacity for cross-presentation translates into a more robust CD8^+^ T cell response to tumor antigens and decreased tumor burden. Taken together, we have identified LAP as a key mechanism that regulates cross-presentation in DCs, emphasizing its importance in maintaining homeostasis and limiting immune activation.

## Results

### *Rubcn^-/-^* DCs display enhanced cross-presentation capacity

To explore the role of LAP in cross-presentation, we generated bone-marrow derived DCs using recombinant Flt3L from both *Rubcn^+/+^* and *Rubcn*^-/-^ mice (Brasel et al., 2000) (Figure S1A-B). On day 6-8, DCs was analyzed by FACS analysis, and cultures over 75% CD11c^+^ CD103^+^ CD8α^+^ were used for *in vitro* experiments. Upon co-culture with apoptotic B16 melanoma cells expressing membrane bound ovalbumin (B16-OVA), *Rubcn^-/-^*DCs had significantly increased levels of the OVA_257-264_ peptide (SIINFEKL) presented in the context of MHC-I (H2-K^b^-OVA_257-264_) in a dose-dependent manner, compared to *Rubcn^+/+^* DCs (Figure 1A). This increase was dependent on phagocytosis, as upregulation of H2-K^b^-OVA_257-264_ expression was abolished when co-culture was performed at 4°C (Figure S1C). Increased expression of H2-K^b^-OVA_257-264_ in *Rubcn^-/-^*DCs was not due to an increase in total H2-K^b^ (MHC-I), as both *Rubcn^+/+^* and *Rubcn^-/-^*DCs increased total H2-K^b^ expression equivalently and in a phagocytosis-dependent manner (Figure 1B, Figure S1C). While studies have demonstrated that DCs deficient for *Atg5* or *Atg7* exhibit reduced recycling of MHC-I molecules, resulting in increased MHC-I antigen presentation (Loi et al., 2016), our data suggests that *Rubcn*-deficiency has no effect on total MHC-I expression, but rather enhances cross-presentation specifically. Transcriptional analysis via RNA-seq revealed that upon co-culture with apoptotic Jurkat cells, *Rubcn^-/-^* DCs upregulated components of cross-presentation (such as *Canx*, *Ciita*, and *Erap1*) and activation machinery (such as *CD40* and *CD86*), more robustly than *Rubcn^+/+^* DCs (Figure 1C). Similar to flow cytometric analysis, *H2k1* expression was equivalent in *Rubcn^+/+^* and *Rubcn*^-/-^ DCs (Figure 1C). We used qPCR to confirm an upregulation of transcriptional activity at 6 hours post-stimulation and observed that *Rubcn^-/-^* DCs displayed increased expression of *Tap1*, *Tap2*, and *B2m*, compared to *Rubcn^+/+^* DCs (Figure S1D). While *Rubcn^+/+^*and *Rubcn^-/-^* DCs had equivalent levels basally, *Rubcn^-/-^* DCs displayed increased surface expression of CD86, a co-stimulatory molecule (Greenwald et al., 2005), in response to co-culture with apoptotic B16-OVA cells (Figure 1D). In addition, and similar to *Rubcn^-/-^*macrophages, *Rubcn^-/-^* DCs significantly upregulated pro-inflammatory cytokines and chemokines, compared to *Rubcn^+/+^* DCs (Martinez et al., 2011a; Martinez et al., 2016b). Specifically, *Rubcn^-/-^* DCs upregulated factors known to promote CD8^+^ T cell recruitment (such as *Ccl2*, *Ccl3*, *Ccl4*, *Cxcl9*, and *Cxcl10*) and activation (such as *Il1b*, *Il6*, *Il12b*, and *Tnf*) (Figure 1C, S1E) (Rahimi and Luster, 2018; Turner et al., 2014). Consistent with previous reports (Hayashi et al., 2018), *Rubcn^-/-^* DCs produced less IFNβ than *Rubcn^+/+^* DCs (Figure 1C, S1E). Therefore, *Rubcn^-/-^* DCs significantly upregulate key mediators known to promote an inflammatory response and enhance CD8^+^ T cell activation.

**Figure 1:**
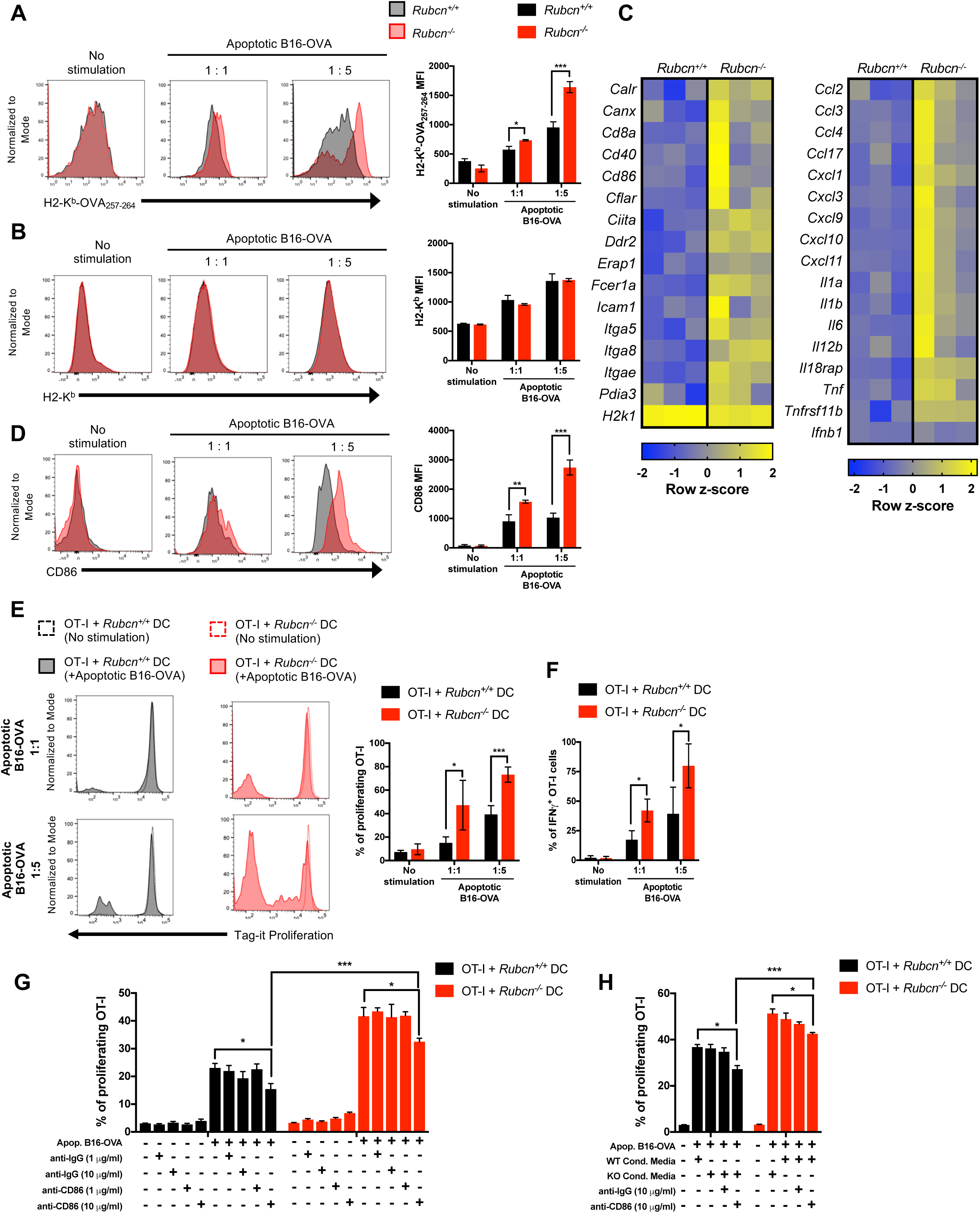
*Rubcn^-/-^* DCs display enhanced cross-presentation capacity. Bone marrow-derived dendritic cells (DCs) were generated from *Rubcn^+/+^* (black) and *Rubcn^-/-^* (red) mice *in vitro* with FLT3-L for 7 days. (**A-B, D**) DCs were co-cultured with apoptotic B16-OVA cells (1 or 5 apoptotic cells: 1 DC). Eighteen hours later, DCs were harvested for flow cytometry analysis of H2-K^b^-OVA_257-264_ (**A**), H2-K^b^ (**B**), and CD86 (**D**) expression. (**C**) DCs were co-cultured with apoptotic Jurkat cells (5 apoptotic cells: 1 DC). Thirty minutes later, RNA was harvested for RNA-seq. Genes with fold change (FC>1.25) compared to no stimulation and p<0.05 (ANOVA) were considered significant and entered in DAVID and R studio for further pathway analysis. Heatmap of differentially regulated antigen presentation, activation, and cytokine/chemokine machinery depicted as row z-score is shown (n=3 per genotype). (**E-H**) DCs were co-cultured with apoptotic B16-OVA cells (1 or 5 apoptotic cells: 1 DC). Eighteen hours later, loaded DCs were co-cultured with naïve OT-I CD8^+^ T cells, labeled with Tag-it Proliferation dye. Forty-eight hours later, *in vitro* cultures were harvested for flow cytometry analysis of OT-I proliferation (**E, G-H**) or IFNγ production (**F**). Prior to OT-I addition, DCs were incubated with anti-IgG, anti-CD86, and/or conditioned media from independent cultures for at least 1 hour (**G-H**), as indicated. Data are expressed as mean ± SEM. No less than three independent experiments were performed, with 3-5 replicates per condition. Significance was calculated using 2-way ANOVA (*p<0.05, **p<0.01, ***p<0.001).

A recent study identified a process akin to LAP, called LC3-associated endocytosis (LANDO), wherein uptake of particles during receptor-mediated endocytosis (RME) results in Rubcn-dependent LC3-decorated, cargo-containing endosomes (Heckmann et al., 2019). Therefore, we also examined the ability to *Rubcn^+/+^* and *Rubcn*^-/-^ DCs to upregulate cross-presentation in response to endocytosis of soluble ovalbumin protein (sOVA protein), a well-characterized *in vitro* model antigen for cross-presentation (Blachere et al., 2005; Burgdorf et al., 2006; Cebrian et al., 2011; Yatim et al., 2015). Similar to apoptotic B16-OVA cell co-culture, endocytosis of sOVA protein induces significantly increased expression of H2-K^b^-OVA_257-264_ by *Rubcn*^-/-^ DCs, compared to *Rubcn^+/+^* DCs (Figure S1F). Importantly, both *Rubcn^+/+^* and *Rubcn*^-/-^ DCs upregulated H2-K^b^-OVA_257-264_ expression at equivalent levels in response to stimulation with OVA_257-264_ peptide, indicating that cross-presentation, but not classical MHC-I antigen presentation, was increased in the absence of *Rubcn* (Figure S1F).

To test their capacity to activate CD8^+^ T cells, *Rubcn^+/+^* and *Rubcn^-/-^* DCs were loaded with apoptotic B16-OVA cells, followed by co-culture with naïve CD8^+^ T cells from OT-I mice, which express a transgenic T cell receptor that recognizes the OVA_257-264_ peptide in the context of MHC-I (Hogquist et al., 1994). OT-I T cells were labeled with Tag-it Proliferation Dye prior to the co-culture, and proliferation and activation were assessed 48 hours later. Co-culture with *Rubcn^+/+^* DCs resulted in a modest amount of proliferation of OT-I T cells. Consistent with higher levels of cross-presentation machinery, co-culture with *Rubcn^-/-^* DCs resulted in a more robust proliferation of OT-I T cells (Figure 1E). OT-I T cells co-cultured with *Rubcn^-/-^* DCs also demonstrated increased effector function, as evidenced by increased IFNγ production (Figure 1F). Similarly, OT-I T cells co-cultured with sOVA protein-loaded *Rubcn^-/-^*DCs also demonstrated increased proliferation and effector function. No difference was in proliferation or activation was observed in OT-I T cells co-cultured with OVA_257-264_ peptide-loaded *Rubcn^+/+^* or *Rubcn^-/-^* DCs (Figure S1G-H).

Both increased CD86 expression and pro-inflammatory cytokine/chemokine production can positively contribute to CD8^+^ T cell activation (Zhang and Bevan, 2011). To determine if increased CD86 expression on *Rubcn^-/-^* DCs was responsible for increased CD8^+^ T cell activation capacity, we performed OT-I : DC co-cultures in the presence of anti-CD86 blocking antibody or isotype control antibody. Isotype control antibody treatment had no effect on OT-I proliferation, while the higher dose (10 μg/ml) of anti-CD86 blocking antibody inhibited OT-I proliferation modestly with both *Rubcn^+/+^* and *Rubcn^-/-^* DCs (Figure 1G). However, OT-I proliferation in culture with *Rubcn^-/-^* DCs with anti-CD86 inhibition was still significantly increased compared OT-I proliferation in culture with wild type DCs with anti-CD86 inhibition (Figure 1G, S2A). To assess the role of pro-inflammatory cytokine/chemokines produced by *Rubcn^-/-^* DCs, we collected culture media from *Rubcn^+/+^*and *Rubcn^-/-^* DCs after 24 hours of culture with apoptotic B16-OVA cells. This conditioned media was then used in new OT-I : DC co-cultures stimulated with apoptotic B16-OVA cells. The presence of *Rubcn^-/-^* (KO) conditioned media in OT-I: *Rubcn^+/+^* DC cultures did not confer a proliferative advantage, and the presence of *Rubcn^+/+^* (WT) conditioned media did not inhibit proliferation of OT-I co-cultured with *Rubcn^-/-^* DCs (Figure 1H, S2B). Additionally, anti-CD86 inhibition had no effect on OT-I proliferation in conditioned media co-cultures (Figure 1H, S2B). Therefore, increased cross-presentation capacity, not increased co-stimulation or inflammatory cytokine/chemokine production, is the main contributing factor to increased antigen presentation capacity of *Rubcn^-/-^* DCs.

Although multiple molecular components overlap between canonical autophagy and LAP (and LANDO), only canonical autophagy requires the activity of the pre-initiation complex, composed of ATG13, FIP200, and ULK1 (Martinez et al., 2011a). We and others have reported that *Rubcn* deficiency can result in increased canonical autophagic activity (Martinez et al., 2015). To confirm that RUBCN’s effect on cross-presentation occurs independently of canonical autophagy, we examined the cross-presentation capacity of *Ulk1^+/+^* and *Ulk1^-/-^* DCs. *Ulk1^+/+^* and *Ulk1^-/-^* DCs displayed equivalent expression levels of H2-K^b^-OVA_257-264_ upon stimulation with either sOVA protein or apoptotic B16-OVA cells (Figure S2C). As such, OT-I T cells co-cultured with either *Ulk1^+/+^* and *Ulk1^-/-^* DCs proliferated similarly (Figure S2D), indicating that defects in LAP, not canonical autophagy, is responsible for increased cross-presentation in *Rubcn^-/-^* DCs. We further confirmed this finding with DCs deficient for *Atg7* from *Cd11c-Cre Atg7^flox/flox^* mice, which contain DCs deficient for both canonical autophagy and LAP (Martinez et al., 2011a; Sanjuan et al., 2009). *Atg7*-deficient DCs also display increased cross-presentation capacity as evidenced by increased expression of H2-K^b^-OVA_257-264_ (Figure S2E) and increased OT-I T cell proliferation in co-culture (Figure S2F). Collectively, these data demonstrate that LAP- and LANDO-deficiency confer increased cross-presentation capacity.

### *Rubcn^-/-^* DCs engulf cargo, yet retain cargo in less degradative phagosomes

To address how the absence of LAP imparts an increased capacity for cross-presentation, we next examined the nature of the cargo-containing phagosome in the absence of *Rubcn*. *Rubcn^+/+^* and *Rubcn^-/-^*DCs phagocytosed apoptotic cells (Figure 2A) equivalently, so we utilized flow cytometry to analyze LC3 localization on cargo-containing phagosomal membranes. Briefly, apoptotic B16-OVA cells were labeled with PKH26 and co-cultured with *Rubcn^+/+^* and *Rubcn^-/-^*DCs that express the transgene for GFP-LC3 (Martinez et al., 2015). After four hours, DCs were harvested, permeabilized on ice with digitonin (20 µg/ml), which disrupts the plasma membrane, but leaves intracellular membranes intact (Martinez et al., 2015). GFP-LC3 translocation to the PKH26^+^ apoptotic cell-containing phagosome was analyzed via flow cytometry by examining mean fluorescent intensity (MFI) of GFP-LC3 associated with PKH26^+^ events (Shvets and Elazar, 2009). While *Rubcn^+/+^*DCs translocated GFP-LC3 to apoptotic cell-containing phagosomes, *Rubcn^-/-^* DCs contained significantly reduced GFP-LC3 association with apoptotic cell-containing phagosomes (Figure 2B).

**Figure 2:**
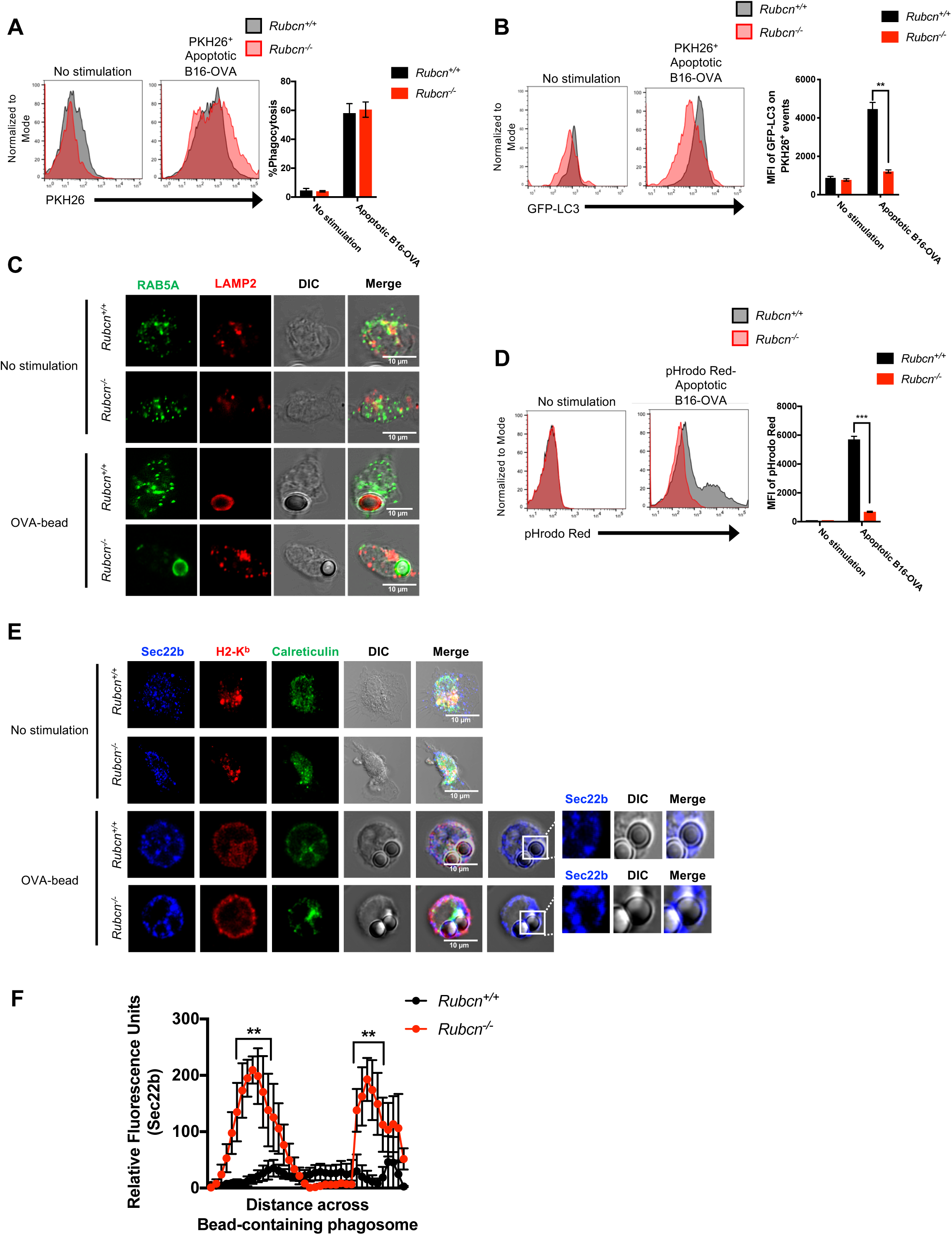
*Rubcn^-/-^* DCs engulf cargo, yet retain cargo in less degradative phagosomes. Bone marrow-derived dendritic cells (DCs) were generated from *Rubcn^+/+^*(black) and *Rubcn^-/-^* (red) mice *in vitro* with FLT3-L for 7 days. (**A-B**) GFP-LC3^+/+^DCs were co-cultured with PKH26-labeled apoptotic B16-OVA cells (5 apoptotic cells: 1 DC). After one hour, phagocytosis of PKH26-labeled apoptotic B16-OVA cells was analyzed by flow cytometry (**A**). After four hours, DCs were harvested, permeabilized by digitonin (20 µg/ml), and GFP-LC3 fluorescence associated with PKH26^+^ events was analyzed by flow cytometry (**B**). (**C**) DCs were co-cultured with OVA-coated latex beads (5 beads: 1 DC) in chamber slides. Four hours later, DCs were fixed, permeabilized, stained for RAB5A (green) and LAMP2 (red), and imaged via confocal microscopy. Representative images are shown. (**D**) Apoptotic B16-OVA cells were labeled with pHrodo Red per manufacturer’s instructions, then co-cultured with *Rubcn^+/+^* and *Rubcn^-/-^* DCs *in vitro*. Four hours later, pHrodo Red fluorescence was measured in DCs by flow cytometry, as a readout of phagosomal maturation. (**E**) DCs were co-cultured with OVA-coated latex beads (5 beads: 1 DC) in chamber slides. Four hours later, DCs were fixed, permeabilized, stained for Sec22b (blue), H2-Kb (red), and Calreticulin (green), and imaged via confocal microscopy. Representative images are shown. (**F**) Signal intensity profile for Sec22b across phagocytosed beads was quantified by Fiji. Data (intensity measurements across beads) are presented as mean ± SEM (n = 10-15 cells per genotype). Flow cytometric data are expressed as mean ± SEM. No less than three independent experiments were performed, with 3-5 replicates per condition. Significance was calculated using 2-way ANOVA (*p<0.05, **p<0.01, ***p<0.001).

Compared to macrophages, DCs are considered more efficient at cross-presentation, as they retain engulfed cargo in less acidified compartments, allowing for longer periods of antigen integrity (Belizaire and Unanue, 2009). To explore the nature of cargo-containing phagosomes in *Rubcn^-/-^* DCs, we utilized latex beads coated with OVA protein and performed immunofluorescence for localization of RAB5A, a marker of early endosomes, and LAMP2, a marker of lysosomes (Henault et al., 2012). Unstimulated *Rubcn^+/+^* and *Rubcn^-/-^* DCs exhibited equivalent RAB5A and LAMP2 staining (Figure 2C), and both genotypes displayed equivalent levels of lysosomal enzymes, including Cathepsin S (*Ctss*) which has been shown to be critical for cross-presentation (Shen et al., 2004), suggesting no inherent lysosomal defect in *Rubcn^-/-^* DCs (Figure S3A). However, LAMP2 robustly translocated to OVA-bead-containing phagosomes in *Rubcn^+/+^* DCs, but failed to localize with OVA-bead-containing phagosomes in *Rubcn^-/-^*DCs (Figure 2C). Rather, RAB5A localized with OVA-bead-containing phagosomes in *Rubcn^-/-^*DCs, indicative of their relative immaturity (Figure 2C). We further examined the nature of cargo-containing phagosomes in *Rubcn^-/-^*DCs using apoptotic B16-OVA cells labeled with pHrodo Red (Jantas et al., 2015), a pH-sensitive dye that increases fluorescence in acidic environments, such as lysosomes. *Rubcn^+/+^* DCs exhibited significant pHrodo Red fluorescence when co-cultured with pHrodo-labeled apoptotic B16-OVA cells, yet *Rubcn^-/-^* DCs displayed minimal pHrodo Red fluorescence upon co-culture, consistent with their decreased LAMP2 localization at the cargo-containing phagosome (Figure 2D). We also observed a decreased capacity for clearance of apoptotic cells *in vitro* by *Rubcn^-/-^* DCs, consistent with retention of engulfed cargo in a less degradative environment (Figure S3B). Similar data were obtained using pHrodo-labeled sOVA protein, wherein *Rubcn^-/-^* DCs displayed minimal pHrodo Red fluorescence upon stimulation, suggesting that both LAP- and LANDO-deficiency results in defects in the maturation of cargo-containing vesicles (Figure S3C).

Recent studies suggest that the active translocation of ERGIC-associated proteins to the phagosome via a Sec22b-mediated mechanism is required for cross-presentation. It is believed that Sec22b functions by both promoting phagosome-to-cytosol escape of the cargo, as well as inhibiting phagosomal-lysosomal maturation (Cebrian et al., 2011; Wu et al., 2017). *Rubcn^+/+^* and *Rubcn^-/-^* DCs displayed equivalent localization of Sec22b with the ERGIC marker, Calreticulin, at a resting state (Figure 2E, Figure S3D). In the absence of *Rubcn*, however, Sec22b co-localization with OVA-bead-containing phagosomes was significantly increased, compared to wild type DCs (Figure 2E-F). Taken together, these data demonstrate that *Rubcn* deficiency results in retention of phagocytosed material in immature phagosomal compartments with increased association of the pro-cross-presentation molecule, Sec22b.

### *Rubcn^-/-^* DCs exhibit increased phagosome-to-cytosol escape and proteasome-mediated generation of peptides

To examine phagosomal escape, we utilized two different approaches. In the first assay (Cebrian et al., 2011; Keller et al., 2013), *Rubcn^+/+^* and *Rubcn^-/-^*DCs were loaded with the cephalosporin derived FRET substrate, CCF4, which accumulates in the cytosol and emits a FRET signal at 535 nm. Apoptotic B16-OVA cells were added to the CCF4-loaded DCs in the absence or presence of β-lactamase at 4°C or 37°C. Upon export to the cytosol, β-lactamase cleaves CCF4, resulting in a reduced 535 nm FRET signal and increased emission at 450 nm (Loi et al., 2016). Flow cytometry was used to measure the ratiometric values between both signals (450 nm : 535 nm, as a ratio of cleaved CCF4 to uncleaved CCF4) as a means of examining β-lactamase export to the cytosol (Loi et al., 2016). As demonstrated by increased ratio values, *Rubcn^-/-^* DCs exhibited increased cleaved CCF4 fluorescent signal (and a decrease in uncleaved CCF4 signal), upon co-culture with apoptotic B16-OVA cells at both 90 and 180 minutes, compared to *Rubcn^+/+^* DCs (Figure 3A-B). This increase in escape was dependent on phagocytosis, as cells cultured at 4°C displayed minimal cleaved CCF4 fluorescent signal (Figure 3A-B).

**Figure 3:**
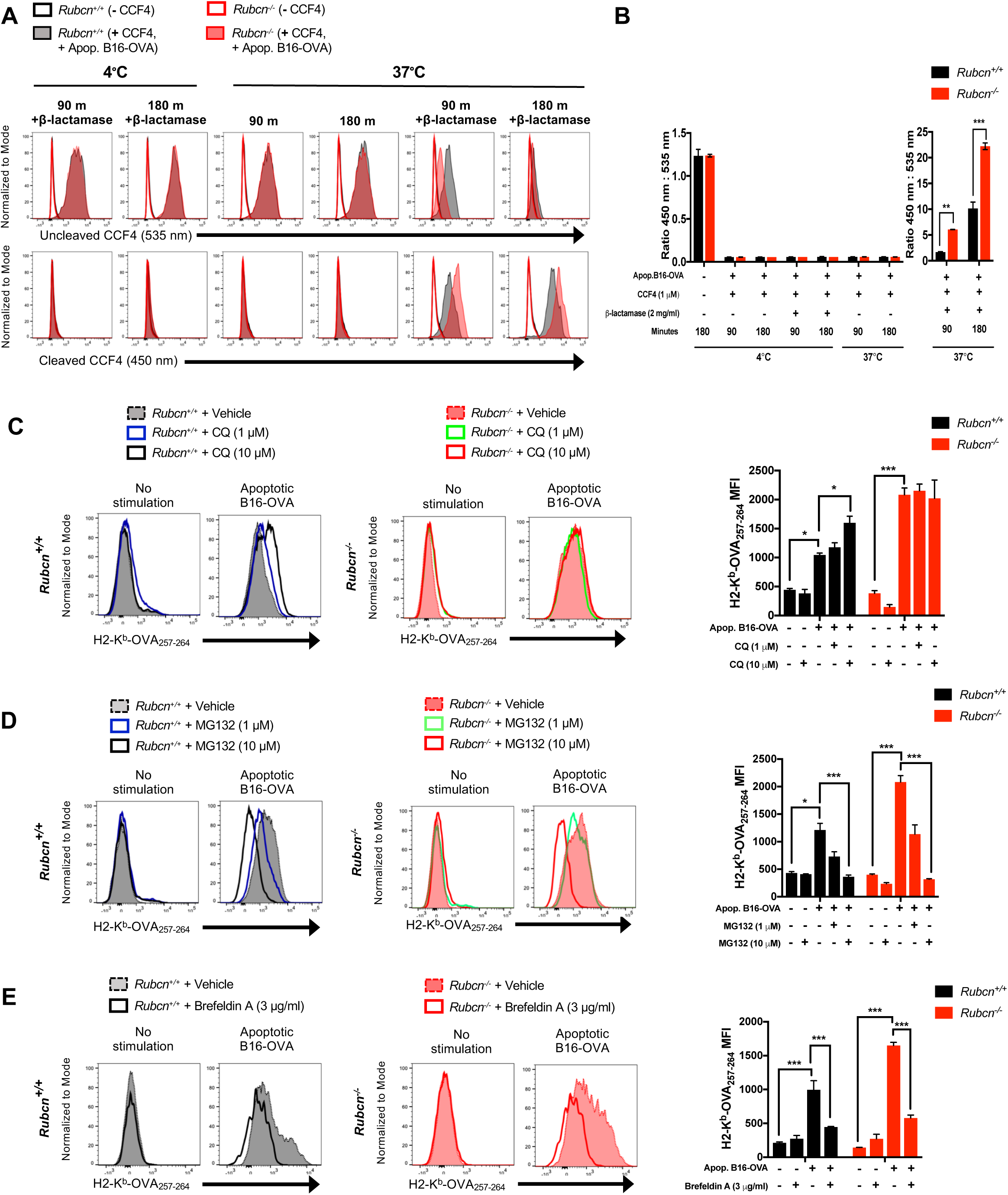
*Rubcn^-/-^* DCs exhibit increased phagosome-to-cytosol escape and proteasome-mediated generation of peptides. Bone marrow-derived dendritic cells (DCs) were generated from *Rubcn^+/+^* (black) and *Rubcn^-/-^* (red) mice *in vitro* with FLT3-L for 7 days. (**A-B**) DCs were loaded with 1 µm CCF4, then co-cultured with apoptotic B16-OVA in the absence or presence of β-lactamase (2 mg/ml) for 90 or 180 minutes at 4°C or 37°C. Uncleaved CCF4 was measured by flow cytometry at an emission of 535 nm, and cleaved CCF4 was measured at an emission of 450 nm. Ratios of 450 nm : 535 nm was calculated by dividing the 450 nm MFI by the 535 nm MFI from a single sample. (**C**) DCs were pre-treated with vehicle or chloroquine (CQ) at 1 or 10 µM for 2 hours and then co-cultured with apoptotic B16-OVA cells (5 apoptotic cells: 1 DC). Eighteen hours later, DCs were harvested for flow cytometry analysis of H2-K^b^-OVA_257-264_ expression. (**D**) DCs were pre-treated with vehicle or MG-132 at 1 or 10 µM for 2 hours and then co-cultured in fresh media with apoptotic B16-OVA cells (5 apoptotic cells: 1 DC). Eighteen hours later, DCs were harvested for flow cytometry analysis of H2-K^b^-OVA_257-264_ expression. (**E**) DCs were pre-treated with vehicle or Brefeldin A at 3 µg/ml for 2 hours and then co-cultured in fresh media with apoptotic B16-OVA cells (5 apoptotic cells: 1 DC). Eighteen hours later, DCs were harvested for flow cytometry analysis of H2-K^b^-OVA_257-264_ expression. Data are expressed as mean ± SEM. No less than two independent experiments were performed, with 3-5 replicates per condition. Significance was calculated using 2-way ANOVA (*p<0.05, **p<0.01, ***p<0.001).

Additionally, we incubated *Rubcn^+/+^*and *Rubcn^-/-^* DCs with PKH26-labeled apoptotic B16-OVA cells, and after 4 hours, DCs were treated with a concentration of digitonin mild enough to permeabilize the plasma membrane but leave phagosomal membranes intact (Martinez et al., 2015; Repnik et al., 2016). To confirm that the mild digitonin dose (20 μg/ml) was not permeabilizing lysosomal membranes, we incubated permeabilized cell extracts with the cathepsin-specific fluorogenic substrate, Z-FR-AMC, which emits fluorescence at 460 nm upon cathepsin-mediated cleavage (Repnik et al., 2016). Permeabilization with 20 μg/ml digitonin did not results in a fluorescent signal, indicative of no cathepsin release from lysosomes (Figure S3E). Permeabilization with a higher dose (200 μg/ml) of digitonin, however, resulted in increased cathepsin-mediated fluorescence in both *Rubcn^+/+^* and *Rubcn^-/-^*DCs, suggesting that increased concentrations of digitonin are capable of permeabilizing the membranes of intracellular structures (Figure S3E). Importantly, *Rubcn^+/+^* and *Rubcn^-/-^* DCs displayed equivalent levels of cathepsin-mediated fluorescence at 200 μg/ml digitonin, and transcriptional analysis revealed no difference in total cathepsin content (Figure S3A, S3E). Compared to *Rubcn^+/+^*DCs, *Rubcn^-/-^* DCs displayed significantly increased phagosomal escape of apoptotic B16-OVA cells, as evidenced by increased PKH26^+^ fluorescence in cytosolic extracts of DCs permeabilized with 20 μg/ml digitonin (Figure S3F). Similar results were obtained when *Rubcn^+/+^* and *Rubcn^-/-^* DCs were cultured with Alexa Fluor 555-conjugated sOVA protein (Figure S3G). Taken together, these data demonstrate the RUBCN limits phagosome-to-cytosol antigen escape during LAP (and LANDO).

We next asked if cargo persisting in immature phagosomal compartments contributed to enhanced phagosomal escape. Chloroquine (CQ) is a lysosomotropic weak base widely used as a pharmacological inhibitor of autophagic flux and LAP (Martinez et al., 2011a; Mauthe et al., 2018). Recent studies demonstrate that while other autophagy inhibitors, such as bafilomycin A, inhibit autophagy by increasing the lysosomal pH of autolysosomes, CQ functions by inhibiting autophagosome and lysosome fusion, without modulating lysosomal acidity (Boya et al., 2003; Mauthe et al., 2018). Pre-treatment of *Rubcn^+/+^*, but not *Rubcn^-/-^*, DCs resulted in increased phagosomal escape of apoptotic B16-OVA cells (Figure S3F) (or sOVA protein, Figure S3G), in a dose-dependent manner, suggesting that retention of cargo in less degradative compartments can promote phagosomal escape. Similarly, *Rubcn^+/+^* DCs pre-treated with CQ exhibited significantly increased capacity for cross-presentation, as evidenced by higher levels of H2-K^b^-OVA_257-264_ expression in response apoptotic B16-OVA cells (Figure 3C), supporting the notion that inhibition of phagosomal-lysosomal fusion promotes cross-presentation. CQ treatment of *Rubcn^-/-^* DCs, however, had no effect on their cross-presentation ability. We observed similar results with sOVA protein-induced LANDO (Figure S3H), suggesting that defects in phagosomal maturation in LAP- or LANDO-deficient cells allows for increased phagosome-to-cytosol escape and hence, increased cross-presentation.

Increased escape of engulfed cargo from the phagosome to the cytosol suggests that antigens are now accessible to proteasome-mediated degradation. While it has been reported that peptide generation can occur in the phagosomal compartments, most data indicate that peptide generation occurs primarily via the proteasome pathway (Kovacsovics-Bankowski and Rock, 1995; Palmowski et al., 2006). To examine the contribution of the proteasome to increased cross-presentation in *Rubcn^-/-^*DCs, we pre-treated *Rubcn^+/+^* and *Rubcn^-/-^* DCs with MG-132, a potent cell-permeable proteasome inhibitor (Burgdorf et al., 2006), followed by co-culture with apoptotic B16-OVA cells. Cross-presentation by both *Rubcn^+/+^*and *Rubcn^-/-^* DCs was significantly impaired by proteasome inhibition by MG-132 in a dose-dependent manner, with a reduction in expression of H2-K^b^-OVA_257-264_ (Figure 3D). Pre-treatment with MG-132 also significantly inhibited cross-presentation induced by sOVA protein in both genotypes (Figure S3I). Transcriptional analysis demonstrated that *Rubcn^+/+^*and *Rubcn^-/-^* DCs had equivalent expression levels of key proteasomal subunits, basally and upon apoptotic cell co-culture (Figure S3J). Collectively, our data reveal that, in the absence of *Rubcn*, prolonged retention of engulfed cargo within immature, Sec22b^+^ phagosomal compartments results in increased translocation of antigens from the phagosome to the cytosol and increased accessibility to degradation by the proteasome.

### Cross-presentation by *Rubcn^-/-^* DCs requires ER/Golgi-mediated activity

Classically, proteasome-generated peptides are loaded onto MHC-I molecules via TAP1/2 on the ER, where they can be further trimmed by ERAP1 to MHC-I-compatible lengths, if needed. The peptide-MHC-I complex (pMHC-I) is then trafficked to the plasma membrane via the Golgi apparatus (Cruz et al., 2017). Evidence exists, however, for TAP1-independent pathway, wherein proteasome-generated peptides re-enter the phagosome and are loaded onto MHC-I within the phagosome (Joffre et al., 2012; Nair-Gupta et al., 2014). Therefore, we examined whether TAP1 loading of peptides at the ER and trafficking of pMHC-I via the Golgi apparatus was required for cross-presentation in *Rubcn^+/+^* and *Rubcn^-/-^* DCs. Brefeldin A disrupts Golgi-ER-mediated protein transport by inhibiting the recruitment and activity of GBF1, a guanine nucleotide exchange factor (GEF) required for ADP-ribosylation factor (ARF)-mediated protein transport between the ER and the Golgi (Garcia-Mata et al., 2003). Brefeldin A has been shown to inhibit multiple trafficking pathways, including cross-presentation (Andrieu et al., 2003; Gutierrez-Martinez et al., 2015). Treatment of *Rubcn^+/+^* and *Rubcn^-/-^* DCs with Brefeldin A drastically reduced cross-presentation in response to apoptotic B16-OVA (Figure 3E) or sOVA protein (Figure S3K), suggesting ER/Golgi-mediated protein transport is required for cross-presentation. While peptide loading onto MHCI molecules can occur post-Golgi (Cruz et al., 2017; Joffre et al., 2012), the ability of Brefeldin A to inhibit cross-presentation, in conjunction with the upregulation of *Erap1* (Figure 1C) and *Tap1* (Figure S1D), suggests that the cytosolic pathway is the predominant process in our system.

### *Cd11c-Cre Rubcn^flox/flox^* mice display decreased tumor burden

The ability to mount an effective anti-tumor response requires cross-presentation of cell-associated antigens *in vivo* to elicit robust CD8^+^ T cell activity (Alloatti et al., 2017). To investigate the role of LAP in cross-presentation, we utilized mice harboring a DC-specific deletion of the *Rubcn* gene (*Cd11c-Cre Rubcn^flox/flox^*). As controls, we used littermates that expressed the floxed *Rubcn* gene but did not express the Cre recombinase (*Rubcn^flox/flox^*) (Figure S4A). B16-OVA cells were injected intradermally into *Rubcn^flox/flox^* and *Cd11c-Cre Rubcn^flox/flox^* mice, and tumor growth and intratumoral immune infiltration and activation were monitored. DC-specific deletion of *Rubcn* resulted in significantly delayed tumor growth (Figure 4A) and significantly less tumor burden (Figure 4B, S4B) *in vivo.* In addition, tumors from *Cd11c-Cre Rubcn^flox/flox^* mice contained more CD45^neg^ Zombie-UV^+^ dead cells, compared to *Rubcn^flox/flox^* mice, suggesting increased tumor cell death (Figure 4C). As we observed an increase in the percentage of CD45^+^ cells in the tumors of *Cd11c-Cre Rubcn^flox/flox^* mice (Figure 4D), we next examined the immune composition of the tumor (Figure S4C). On day 16, we observed that tumors from *Cd11c-Cre Rubcn^flox/flox^* mice contained more CD11c^+^ CD103^+^ CD8α^+^ cross-presenting DCs than tumors from *Rubcn^flox/flox^* mice (Figure 4E). These *Rubcn*-deficient DCs also expressed significantly increased levels of the tumor-derived OVA_257-264_ peptide (SIINFEKL) presented in the context of MHC-I (H2-K^b^-OVA_257-264_) (Figure 4F). No differences were detected in percentage or activation status of tumor-associated macrophages from *Rubcn^flox/flox^* and *Cd11c-Cre Rubcn^flox/flox^* mice (Figure S4D). Moreover, the dLNs from *Cd11c-Cre Rubcn^flox/flox^* mice contained more cross-presenting DCs, compared to *Rubcn^flox/flox^*mice (Figure 4G).

**Figure 4:**
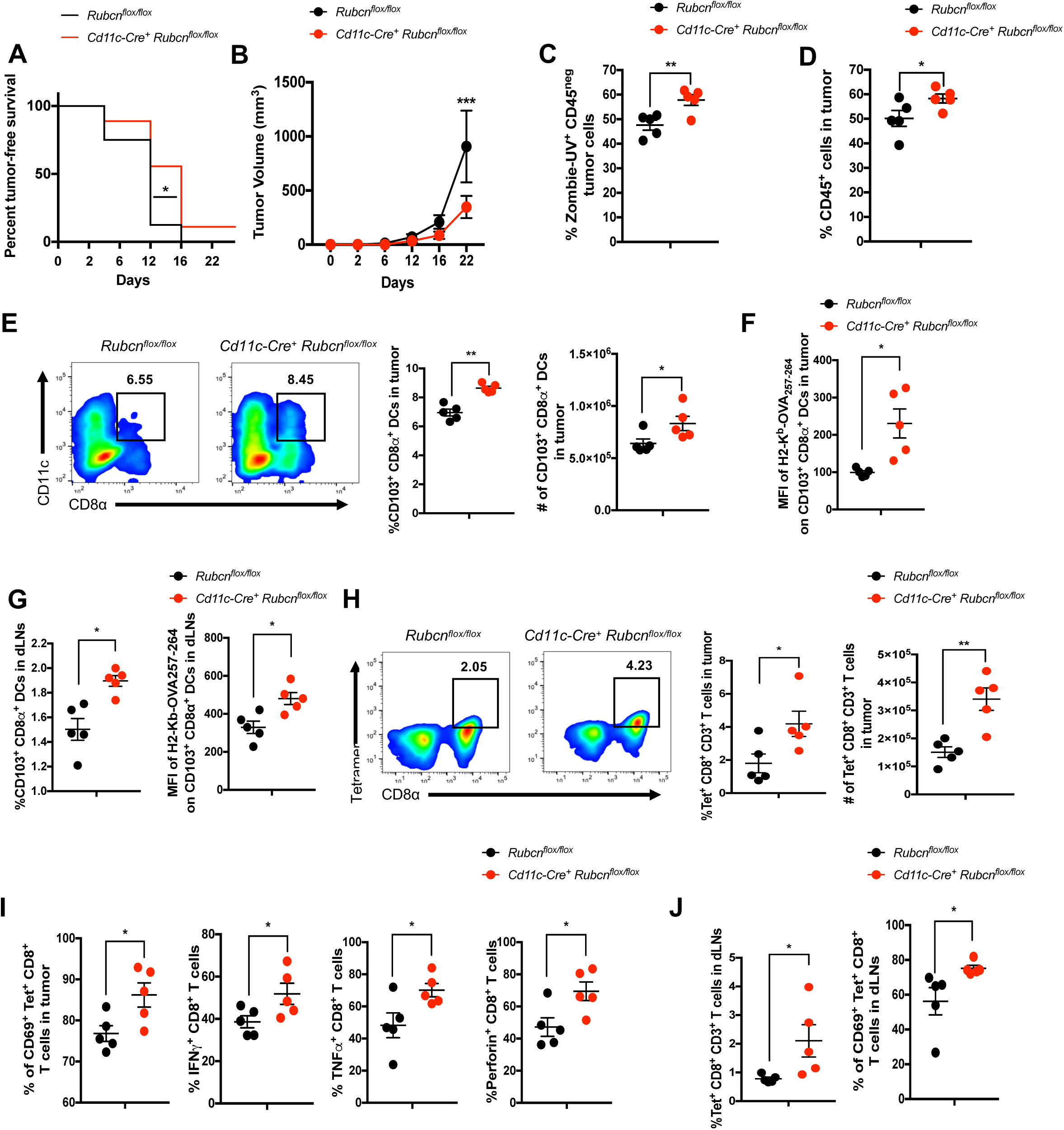
*Cd11-Cre Rubcn^flox/flox^* mice display decreased tumor burden and increased antigen-specific CD8^+^ T cell activation. 1 x 10^6^ B16-OVA cells were injected intradermally into *Rubcn^flox/flox^*(black, n=5) or *Cd11-Cre^+^ Rubcn^flox/flox^* (red, n=5) mice. (**A-B**) Tumor measurements were recorded every 2-4 days. Significance for **A** was calculated using Mantel-Cox Test, and significance for **B** was calculated using 2-way ANOVA (*p<0.05, **p<0.01, ***p<0.001). (**C-E**) On day 16 post-injection, tumors were harvested for flow cytometry analysis. Tumors were assessed for tumor cell death (Zombie-UV^+^ CD45^neg^, **C**) and immune cell infiltration (CD45^+^, **D**). (**E-F**) Representative flow plots for CD11c^+^ CD103^+^ CD8α^+^ DCs. Percentage of and number of CD11c^+^ CD103^+^ CD8α^+^ DCs within CD45^+^ population (**E**); MFI of H2-K^b^-OVA_257-264_ expression (**D**) are expressed as mean ± SEM. (**G**) On day 16 post-injection, dLNs were harvested for flow cytometry analysis. Percentage of CD11c^+^ CD103^+^ CD8α^+^ DCs and MFI of H2-K^b^-OVA_257-264_ expression are expressed as mean ± SEM. (**H-I**) On day 16 post-injection, tumors were harvested for flow cytometry analysis. Representative flow plots for CD8α^+^ Tetramer^+^ T cells are shown in **G**. Percentage of and number of CD8α^+^ Tetramer^+^ T cells within CD45^+^ population (**H**) and percentage of CD69^+^, IFNγ^+^, TNFα^+^, and Perforin^+^ CD8+ T cells within tumor (**I**) are expressed as mean ± SEM. (**J**) On day 16 post-injection, dLNs were harvested for flow cytometry analysis. Percentage of CD8α^+^ Tetramer^+^ T cells and percentage of CD69^+^ CD8+ T cells are expressed as mean ± SEM. No less than three independent experiments were performed, with 5 replicates per genotype. Significance was calculated using 2-way ANOVA (*p<0.05, **p<0.01, ***p<0.001).

We then examined the endogenous OVA-specific CD8^+^ T cell response using MHC-I peptide tetramers of SIINFEKL, referred hereafter as Tetramer (Snyder et al., 2019; Yatim et al., 2015). Tumors from *Cd11c-Cre Rubcn^flox/flox^* mice contained significantly more Tetramer^+^ CD8^+^ T cells, indicating that *Rubcn*-deficient DCs are more efficient at cross-presentation of tumor-derived antigens (Figure 4H). These OVA-specific CD8^+^ T cells were more activated, as evidenced by increased expression of CD69, and more functional, as evidenced by increased production of IFNγ, TNFα, and Perforin (Figure 4I). Similarly, dLNs from *Cd11c-Cre Rubcn^flox/flox^* mice also contained more Tetramer^+^ CD8^+^ T cells that displayed a more activated phenotype, compared to dLNs from *Rubcn^flox/flox^*mice (Figure 4J). These trends continued, wherein tumors from *Cd11c-Cre Rubcn^flox/flox^* mice contained DCs with increased expression of H2-K^b^-OVA_257-264_ (Figure S4E) and more activated Tetramer^+^ CD8^+^ T cells (Figure S4F) at day 22. Overall, these studies demonstrate that *Cd11c-Cre Rubcn^flox/flox^*mice are able to cross-present tumor antigens more efficiently and mount a more robust tumor-specific CD8^+^ T cell response, resulting in reduced tumor burden.

## Discussion

The most abundant source of antigen for cross-presentation is the host, as host cells undergo a continuous cycle of death and homeostatic clearance. In this regard, the MHC-I pathway serves as necessary quality control of the state of self, facilitating the presentation of both endogenous proteins and antigens derived from exogenously acquired dying cells during efferocytosis. Whereas presentation of peptides generated from healthy cells elicits only a modest CD8^+^ T cell response (as these highly autoreactive T cells undergo negative selection in the thymus), presentation of peptides from cells undergoing stress, such as transformation or viral infection, can result in a robust CD8^+^ T cell response (Cruz et al., 2017). Efficient activation of CD8^+^ T cells via cross-presentation, however, can be a double-edged sword. Defects in cross-presentation results in decreased anti-tumor responses and increased tumor burden (Alloatti et al., 2017). Conversely, heightened cross-presentation is associated with increased autoimmunity, as self-antigens are presented to and activate CD8^+^ T cells more readily (Calderon and Unanue, 2012; Ji et al., 2013). Thus, control of the cross-presentation pathway is a critical node in preserving the balance between homeostasis and inflammation.

Here, we demonstrate that a form of non-canonical autophagy, LAP, and its related process, LANDO, function to restrict cross-presentation in DCs. Mechanistically, we describe that LAP functions as a negative regulator of cross-presentation by promoting the maturation of the cargo-containing phagosome to limit phagosome-to-cytosol escape and antigen access to the proteasome. In the absence of *Rubcn*, which is required for LAP but not canonical autophagy, phagosomal maturation is significantly blunted, as LC3-II fails to decorate the phagosome to promote fusion with the lysosome. Within *Rubcn^-/-^* DCs, cargo persists in immature vesicles, which also display increased association of Sec22b, a pro-cross-presentation molecule. This results in increased phagosome-to-cytosol antigen escape, subsequent processing of antigen into peptides, and heightened presentation of pMHC-I on the plasma membrane via ER/Golgi activity.

Recent studies have demonstrated a role for ATG5 in MHC-II-mediated antigen presentation during experimental autoimmune encephalomyelitis, wherein CD4^+^ T cell activation was reduced in *Cd11c-Cre^+^ Atg5^flox/flox^*mice (Keller et al., 2017). This suggests that LAP is required for MHC-II antigen presentation, though studies examining *Rubcn’s* role in MHC-II antigen presentation are needed. While other studies have implicated macroautophagy (via ATG5 and ATG7) in the recycling of MHC-I molecules (Loi et al., 2016), we did not observe an increase in total MHC-I (H2-K^b^) expression in *Rubcn^-/-^* DCs at the transcriptional or protein level. As RUBCN can also inhibit canonical autophagy (Martinez et al., 2015), it is possible that increased autophagy conferred by the absence of *Rubcn* accounts for this difference in observations, suggesting that further studies detailing the crosstalk between these two related pathways during cross-presentation are needed.

Phagosomes from *Rubcn^-/-^* DCs exhibit increased Sec22b association, which serves two pro-cross-presentation purposes. Firstly, Sec22b plays a role in inhibiting phagosomal maturation. Secondly, Sec22b also promotes escape of engulfed cargo from the phagosome to the cytosol, where it is vulnerable to proteasome-mediated processing into peptides (Alloatti et al., 2017; Cebrian et al., 2011). The mechanisms by which RUBCN regulates Sec22b association with the phagosome is an outstanding question, though. It is predicted that Sec22b can directly interact with RAB5A, which is a classical marker of endosomes and immature phagosomes (Rouillard et al., 2016). Our data demonstrate that engulfed cargo persists in RAB5A^+^ phagosomes in the absence of RUBCN, suggesting that Sec22b co-localization could be due to increased RAB5A presence. Therefore, our data demonstrate that RUBCN employs multiple strategies to limit cross-presentation and subsequent autoreactivity.

Selective inhibition of endosome-associated TAP1 results in inhibition of cross-presentation induced by high doses of sOVA protein, suggesting that ER/Golgi-mediated trafficking of pMHC-I is not required for cross-presentation in this scenario (Burgdorf et al., 2008). However, this study also demonstrated that TLR4/MyD88-mediated translocation of TAP1 to endosomes was not required for efficient cross-presentation of sOVA protein at a dose (200 μg/ml) roughly equivalent to the dose used in our study (150 μg/ml), suggesting that cargo can enter the cross-presentation pathway at different points (Burgdorf et al., 2008). Inhibiting ERAP1, the ER enzyme responsible for trimming peptides into an MHC-I suitable form, or IRAP1, its endosomally localized relative, each reduces cross-presentation by approximately half, with silencing both conferring an additive effect (Saveanu et al., 2009). As *Rubcn^-/-^* DCs demonstrated increased expression of *Tap1* and *Erap1* (but not in *Irap1*), as well as exhibited decreased cross-presentation capacity in response to Brefeldin A treatment, we hypothesize that *Rubcn*-deficiency promotes cross-presentation via TAP1-dependent, ER/Golgi-mediated activity. Further studies using tools that inhibit TAP1 at the ER or phagosome are required to further define these molecular mechanisms.

In addition to the uptake of dying cells, viruses can trigger cross-presentation. While some viruses, such as human immunodeficiency virus (HIV), are cross-presented in a proteasome- and TAP-dependent manner (Maranon et al., 2004), other viruses, such as influenza A virus (IAV) and hepatitis B virus (HBV), have been reported to utilize the vacuolar pathway (Stober et al., 2002). Engagement of TLR signaling during cross-presentation is required for translocation of MHC-I to cargo-containing phagosomes, where vacuolar cross-presentation can occur (Nair-Gupta et al., 2014). Whether RUBCN plays a role in virus-initiated cross-presentation or the vacuolar pathway of cross-presentation remains to be determined.

Increased cross-presentation can also be a contributing factor to the age-related autoimmunity and inflammation observed in *Rubcn^-/-^* mice (Martinez et al., 2016b). Studies have linked cross-presentation to increased autoimmunity. NOD mice, which are susceptible to the development of diabetes mellitus, fail to develop invasive intra-islet insulitis or become diabetic in the absence of MHC-I (Marino et al., 2012; Serreze et al., 1997). Compared to wild type mice, however, aged *Rubcn^-/-^* mice exhibit increased levels of activated CD44^+^ CD62L^-^ CD8^+^ T cells in the periphery, suggesting that increased MHC-I-mediated T cell activation is present in the absence of LAP (Martinez et al., 2016b). It is possible that *Rubcn* deficiency bestows improved cross-presentation capacity in other phagocytes, including macrophages, which have been described as poor cross-presenters (Savina and Amigorena, 2007). Thus, tissue or cell type-specific *Rubcn*-deficient animals may exhibit moderate or organ-restricted autoimmune pathology.

Similar to studies utilizing myeloid cell-specific deletion of *Rubcn* (Cunha et al., 2018), mice with *Rubcn*-deficient DCs demonstrate a significant decrease in tumor burden in a xenograft model of tumorigenesis, due to significantly increased cross-presentation capacity and an enhanced tumor antigen-specific CD8^+^ T cell response. As cross-presentation by DCs is required for the effectiveness of anti-PD-1 and anti-CD137 immunotherapies in the B16 tumor model (Sanchez-Paulete et al., 2016), a combination of RUBCN inhibition and immune checkpoint inhibitors, such anti-PD-1 and anti-CTLA-4, may synergize for a more potent anti-tumor CD8^+^ T cell response. If modulation of RUBCN activity affects the expression of PD-L1 or other inhibitory molecules on APCs remains an open avenue to explore.

Due to their diversity, plasticity, and efficient cross-presentation capacity, DCs can be exploited as vaccines to yield improved therapeutics (Saxena and Bhardwaj, 2018). Indeed, DC vaccines, such as Sipuleucel-T, have demonstrated remarkable efficacy in the treatment of prostate cancer via the capacity of autologous DCs to mount an effective and tumor-antigen-specific CD8^+^ T cell response (Kantoff et al., 2010). Downregulation of *Rubcn* expression or activity in DCs *ex vivo* could provide additional benefit and further boost their efficacy *in vivo*. Collectively, our study demonstrates that LAP promotes an immunotolerant response by limiting cross-presentation, potentially as a mechanism to suppress excessive activation of CD8^+^ T cells.

## Acknowledgments

The authors thank the animal husbandry staff at NIEHS, as well as Carl Bortner, Maria Sifre, and Kevin S. Katen of the Flow Cytometry Core (NIEHS). We would also like to thank Seddon Y. Thomas, Hideki Nakano, Thomas H. Oguin, III, Donald N. Cook, and Michael Fessler for thoughtful insights and discussions. This work was supported by the NIH Intramural Research Program 1ZIAES10328601 to J.M.

## Author Contributions

The project was conceived by P.S. and J.M. Mouse experiments were conducted by P.S., F.Z., J.P.K., and S.W.W. G.M., L.D., C.J.T., and E.S. provided technical assistance. S.G. provided bioinformatics support. A.S. and A.O. provided reagents and technical expertise. Writing and editing by P.S., J.P.K., G.M., and J.M.

## STAR Methods

### Lead Contact and Materials Availability

Reagents and mice used in this study will be made available under a standard material transfer agreement. Requests for resources and reagents should be directed to and will be fulfilled by the Lead Contact, Jennifer Martinez.

### Experimental Model and Subject Details

#### Mice

##### Strains

Age (8-12 weeks), gender, and weight-matched littermates were used as wild type controls. Mice were housed and bred in a specific-pathogen-free animal facility at the NIEHS/NIH, under the guidance of the NIEHS Animal Care and Use Committee (ACUC) in accordance with the Guide for the Care and Use of Animals.

*Rubcn^-/-^* mice have been previously described (Martinez et al., 2015). Conditional null (“Flox”) Rubicon mice were made from embryonic stem cells purchased from the European Conditional Mouse Mutagenesis Program (EUCOMM Project 76467). *Rubcn^flox/flox^*transgenic mouse strains were generated at the NIEHS animal facility and bred to *B6.Cg-Tg(Itgax-cre)1-1Reiz/J* (*CD11c-Cre^+^*B6, Jax stock no. 008068) mice to generate *CD11c-Cre^+^ Rubcn^flox/flox^*.

*Ulk1^−/−^*mice were provided by M. Kundu (St. Jude Children’s Research Hospital, Memphis, TN, USA) (Kundu et al., 2008). *Atg7^flox/flox^* mice were obtained from M. Komatsu (Tokyo Metropolitan Institute of Medical Science, Tokyo, Japan) (Komatsu et al., 2005) and bred to *CD11c-Cre^+^* mice to generate *CD11c-Cre^+^ Atg7^flox/flox^*. *C57BL/6-Tg(TcraTcrb)1100Mjb/J* (OT-1 B6, Jax stock no. 003831) mice were purchased from Jackson Laboratories (Bar Harbor, ME, USA) and were used as a source for OVA-specific CD8^+^ T cells (described below). All mouse strains were maintained on a C57BL/6 background.

##### Xenograft tumor model

1×10^6^ B16-OVA cells in endotoxin-free PBS (Thermo Fisher Scientific, Waltham, Massachusetts, USA) were injected intradermally into the shaved back of the female mice. The mice were monitored for tumor growth, and the tumors were measured using digital calipers (Mitutoyo Corporation, Kanagawa, Japan). The tumor volume was determined using the formula, *volume = short axis^2^ × long axis × 0.523* (in mm^3^). The solid tumors and draining lymph nodes (dLNs) were harvested in complete media (DMEM+ 10% FCS+ 1% Penicillin-streptomycin) in a non-treated 6-well plate (Thermo Fisher Scientific, Waltham, Massachusetts, USA). The tumor mass was mechanically dissociated using a sharp blade, and they were then enzymatically dissociated by incubating in DMEM containing 50% Accutase (Life Technologies Corporation, Waltham, Massachusetts, USA), 1% penicillin-streptomycin and 5% collagenase/ hyaluronidase (STEMCELL Technologies Inc., Vancouver, British Columbia, Canada) for 1 hour at 37 °C. The tumor cells were filtered using 70 µm filters and washed with PBS. The single cell suspension was prepared by resuspended the cells in MACs buffer (PBS with 0.5% BSA and 2mM EDTA). The cells were then finally resuspended in 80% Percoll (GE Healthcare Bio-Sciences, Pittsburgh, PA, USA) and carefully layered the 40% Percoll. Once the Percoll gradient was prepared, the cells were centrifuged at 1200 x g with no brakes at 4 °C. Finally, tumor-infiltrating immune cells were collected at the interphase of 80%-40% Percoll gradient. The cells were washed twice with endotoxin-free PBS (400 x g, 5 mins, 4 °C) and finally resuspended in MACs buffer. The dLNs were mashed in complete media, filtered using 70 µm filters, and washed twice with endotoxin-free PBS. The single cell suspension was prepared by resuspended the cells in FACs buffer. Tumor-infiltrating immune cells and dLN cells were used to perform subsequent analysis. All injected cells were subjected to and passed the NIEHS Quality Assurance Laboratory (QAL) testing prior to injection.

#### Cell culture

##### Bone marrow-derived dendritic cells (DC) differentiation

Bone marrow was extracted from the 8-12-week-old male mice. The hind legs were removed and placed in cold complete media. The muscle and connective tissues associated with the femur and tibia were removed. The bones were then flushed with cold RPMI complete media (supplemented with 10% FCS (Hyclone, GE Healthcare Bio-Sciences, Pittsburgh, PA, USA), 1% HEPES (Sigma-Aldrich Corporation, St. Louis, Missouri, United States), 1% Glutamax (Life Technologies Corporation, Waltham, Massachusetts, USA), 1% penicillin-streptomycin (Life Technologies Corporation, Waltham, Massachusetts, USA) and 50 µM tissue culture-grade β-mercaptoethanol (β-ME) (Life Technologies Corporation, Waltham, Massachusetts, USA) until they turned white. The BM cells were then passed through a 70 µm cell strainer. The cells were then centrifuged and treated with 9 ml of ammonium chloride solution (STEMCELL Technologies Inc., Vancouver, British Columbia, Canada) for 10 minutes on ice to lyse the red blood cells (RBCs). The cells were centrifuged twice with endotoxin-free PBS (400 x g, 5 mins, 4 °C) and counted. Freshly isolated BM cells were cultured in complete medium supplemented with 200 ng/mL of recombinant FMS-like tyrosine kinase-3 ligand (rFlt3L) (NIEHS Protein Expression Core Facility) at the density of 2 x 10^6^ viable cells per ml in Nunc™ non-treated flasks (Thermo Fisher Scientific, Waltham, Massachusetts, USA) at 37 °C. The media was exchanged every 2-3 days for fresh complete media with 200 ng/ml rFlt3L.

On day 6-8, Flt3L-derived DCs were harvested by collecting the non-adherent DCs in a centrifuge tube. The adherent DCs were treated with 2mM EDTA (Invitrogen, Carlsbad, CA, USA) for 5 mins to harvest them and were then collected in a centrifuge tube. The DCs were washed with endotoxin-free PBS twice and plated for subsequent *in vitro* experiments. Purity was assessed by FACS analysis, and cultures over 75% CD11c^+^ CD103^+^ CD8α^+^ were used for *in vitro* experiments.

##### DC co-culture

DCs were played at 1 x 10^6^/ml in triplicate in 96 well round bottom plates for *in vitro* experiments. Apoptotic cells (below) were used at a ratio of 1 apoptotic cell : 1 DC or 5 apoptotic cells : 1 DC. Low endotoxin whole soluble OVA (sOVA) protein (Worthington Biochemical, Lakewood, NJ USA) was used at 150 μg/ml. Alexa Fluor 555-conjugated OVA was obtained from Thermo-Fisher Scientific (Waltham, MA, USA) and used at 150 μg/ml. OVA_257-264_ peptide (Invivogen, San Diego, CA, USA) was used at 50 ng/ml. OVA (150 μg/ml) was coupled to 3-micron polystyrene beads (Polysciences Inc., Warrington, PA USA) per manufacturer’s instructions. Chloroquine (CQ) was used at 1 or 10 μM to inhibit lysosomal fusion (Sigma-Aldrich, St. Louis, MO USA). MG-132 was used at 1 or 10 μM to inhibit proteasome activity (Sigma-Aldrich, St. Louis, MO USA). Brefeldin A was used at 3 µg/ml to inhibit protein transport (Thermo-Fisher Scientific, Waltham, MA, USA).

##### Cell lines

B16-OVA cells were obtained from A. Oberst at the University of Washington, Seattle, WA USA. The cells were cultured with complete DMEM media supplemented with 10% heat-inactivated fetal bovine serum (HI-FCS) (Hyclone, GE Healthcare Bio-Sciences, Pittsburgh, PA, USA), 1% penicillin-streptomycin (Penicillin-streptomycin) (Life Technologies Corporation, Waltham, Massachusetts, USA) and 500 µg/ml geneticin (Life Technologies Corporation, Waltham, Massachusetts, USA). The media was changed every 2 days. The cells were harvested by trypsinization and washed twice with endotoxin-free PBS before use. The cells were confirmed as negative for mycoplasma contamination by the NIEHS Quality Assurance Laboratory (QAL). Jurkat cells were obtained from M. Fessler at the NIEHS, RTP, NC USA and cultured with complete RPMI media supplemented with 10% heat-inactivated fetal bovine serum (HI-FCS) (Hyclone, GE Healthcare Bio-Sciences, Pittsburgh, PA, USA) and 1% Penicillin-streptomycin (Life Technologies Corporation, Waltham, Massachusetts, USA). The media was exchanged every 2-3 days for fresh complete media. The cells were harvested by collecting all the non-adherent cells, centrifuging, and washing with endotoxin-free PBS. The cells were confirmed as negative for mycoplasma contamination by the NIEHS Quality Assurance Laboratory (QAL).

##### Induction of Apoptosis in B16-OVA cells and Jurkat cells

The cells were washed with endotoxin-free PBS 2x to remove the media and then UV treated for 20 seconds with 20 J/m^2^ in a UV crosslinker (Fisher Scientific, Hampton, NH, USA) (Martinez et al., 2011b). The cells were then returned to the 37 °C incubator. 16 hours of post-UV treatment, the apoptotic cells were detected by Annexin V and PI staining and performing FACs analysis (Martinez et al., 2016a, b).

##### Naïve CD8^+^ T cells isolation

The spleens from naïve OT1 mice C57BL/6-Tg(TcraTcrb)1100Mjb/J (OT-1 B6, Jax Stock No. 003831 Jackson Laboratories Bar Harbor, ME, USA) were harvested and the CD8^+^ T cells were extracted using EasySep™ Mouse Pan-Naïve T Cell Isolation Kit per manufacturer’s instructions. (STEMCELL Technologies Inc., Vancouver, British Columbia, Canada). Isolated T cells were incubated in serum-free media for 2 hours to synchronize their cell cycles. Cells were then stained with Tag it ™ dye (Biolegend, San Diego, CA, USA) per manufacturer’s instructions.

##### T cell : DC co-culture

CD11c^+^ DCs were isolated from day 7 bone marrow-dervied DCs (above) using EasySep™ manufacturer’s guidelines from the Mouse Pan-DC Enrichment Kit (STEMCELL Technologies Inc., Vancouver, British Columbia, Canada). 1x 10^5^ CD11c^+^ DCs were then co-cultured with apoptotic B16-OVA cells (1 or 5 apoptotic cells : 1 DC), sOVA protein (150 µg/ml), or OVA_257-264_ peptide (50 ng/ml) in triplicate for 18 hours in a 96 well plate. Unstimulated or loaded CD11c^+^ DCs were co-cultured with naïve isolated 1x 10^6^ cell cycle-synchronized, Tag it ™-labeled CD8^+^ OT-I T cells (above) for 2 days (10 T cells : 1 DC). T cell proliferation and production of IFNγ was measured by FACs after 48 hours of co-culture.

##### CD86 inhibition *in vitro*

T cell : DC co-culture was performed as described above with the following modifications. Prior to addition of OT-I T cells, antigen-loaded DCs were incubated with IgG2a isotype control antibody or anti-CD86 blocking antibody (Biolegend, San Diego, CA, USA) at 1 or 10 μg/ml for at least 1 hour. DCs were then co-cultured with naïve isolated 1x 10^6^ cell cycle-synchronized, Tag it ™-labeled CD8^+^ OT-I T cells (above) for 2 days (10 T cells : 1 DC). T cell proliferation was measured by FACs after 48 hours of co-culture.

##### Conditioned media

T cell : DC co-culture was performed as described above with the following modifications. Prior to addition of OT-I T cells, media was removed from antigen-loaded DCs, and DCs were gently washed with PBS. Culture media was collected from independent cultures of *Rubcn^+/+^* and *Rubcn^-/-^*DCs after 18 hours of culture with apoptotic B16-OVA cells. Conditioned media from *Rubcn^-/-^*DCs (KO conditioned media) was transferred to *Rubcn^+/+^* DCs, and conditioned media from *Rubcn^+/+^* DCs (WT conditioned media) was transferred to *Rubcn^-/-^* DCs. Cultures were incubated for at least 1 hour with conditioned media, prior to addition of OT-I T cells. DCs were then co-cultured with naïve isolated 1x 10^6^ cell cycle-synchronized, Tag it ™-labeled CD8^+^ OT-I T cells (above) for 2 days (10 T cells : 1 DC). T cell proliferation was measured by FACs after 48 hours of co-culture.

### Method Details

#### Quantification of phagocytosis

Phagocytosis was calculated using flow cytometry analysis as described in (Martinez et al., 2015). The percentage of phagocytosis equals the number of dendritic cells that have been engulfed PKH26-stained B16-OVA. Data represent a minimum of three independent experiments in which technical triplicates of 50,000 cells per sample were acquired using a Fortessa cytometer (BD, Franklin Lake, NJ, USA).

#### Quantification of GFP-LC3 translocation to the phagosome

GFP-LC3 association with the cargo-containing phagosome was assessed by flow cytometry using methods previously described (Martinez et al., 2015) (Shvets and Elazar, 2009). Briefly, apoptotic B16-OVA cells were labeled with PKH26and co-cultured with *Rubcn^+/+^* and *Rubcn^-/-^*DCs that express the transgene for GFP-LC3. After four hours, GFP-LC3^+^ DCs were harvested, washed once with FACS buffer, and permeabilized on ice with digitonin (20 µg/ml) for 15 minutes. GFP-LC3 association with PKH26^+^ apoptotic cell-containing phagosomes was measured by examining mean fluorescent intensity (MFI) of GFP-LC3 of PKH26^+^ events (Shvets and Elazar, 2009). Data represent a minimum of three independent experiments in which technical triplicates of 50,000 cells per sample were acquired using a Fortessa cytometer (BD, Franklin Lakes, New Jersey, USA).

#### pHrodo labeling

Low endotoxin OVA protein was labeled with pHrodo Red Microscale Labeling Kit (Thermo-Fisher Scientific, Waltham, Massachusetts, USA) per manufacturer’s instructions. B16-OVA cells were labeled with pHrodo Red AM Intracellular pH Indicator kit (Thermo-Fisher Scientific, Waltham, Massachusetts, USA) per manufacturer’s instructions.

#### Cell labeling with PKH26 dye

The apoptotic cells were stained using the PKH26 Red Fluorescent Cell Linker Kits (Sigma-Aldrich, St. Louis, Missouri, USA) per manufacturer’s protocol (Martinez et al., 2016a, b). The cell pellet was resuspended in 10 ml of complete medium, transfer to a fresh sterile conical polypropylene tube, centrifuge at 400 x g for 5 minutes and washed the cell pellet 2 more times with 10 ml of complete medium to ensure removal of unbound dye.

#### Digitonin-based measurement phagosome-to-cytosol escape assay

Phagosomal escape using a modified version of a previously described protocol (Repnik et al., 2016). Briefly, cells were cultured with PKH26-labeled apoptotic B16-OVA cells (5 apoptotic cells: 1 DC) or Alexa Fluor 555-labeled OVA protein (150 µg/ml). Six hours later, DCs were permeabilized with digitonin (20 µg/ml) incubating the plate for 10 minutes on ice with shaking. Membrane portions of the sample were pelleted by centrifugation (14,000 rpm, 1 hour). Cytosolic supernatant portions were collected and analyzed for fluorescence as indicated using Tecan M200 Infiniti Pro microplate reader (Tecan, Männedorf, Switzerland).

For measurement of cytosolic cathepsin, DCs were permeabilized with 20 µg/ml or 200 µg/ml for 10 minutes on ice with shaking. DCs then treated with 5 mM DTT and 30 µM cathepsin-specific fluorogenic substrate, Z-FR-AMC (Sigma-Aldrich, St. Louis, MO USA) at 37°C for 20 minutes. Membrane portions of the sample were pelleted by centrifugation (14,000 rpm, 1 hour). Cytosolic supernatant portions were collected and analyzed for AMC (ex 370 nm; em 460 nm) fluorescence using Tecan M200 Infiniti Pro microplate reader.

#### β-lactamase-based measurement of phagosome-to-cytosol escape assay

Phagosomal escape using a modified version of a previously described protocol (Cebrian et al., 2011; Keller et al., 2013). DCs were loaded with 1 µM FRET substrate, CCF4 (Thermo-Fisher Scientific, Waltham, Massachusetts, US), which accumulates in the cytosol and emits a FRET signal at 535 nm when it is uncleaved for 1 hour at room temperature in EM buffer (120mM NaCl, 7mM KCl, 1.8mM CaCl2, 0.8mM MgCl2, 5mM glucose and 25mM HEPES at pH 7.3). Cells were washed once and co-culture apoptotic B16-OVA cells (5 apoptotic cells : 1 DC) in the absence or presence of 2 mg/ml of purified β-lactamase (Sigma-Aldrich, St. Louis, MO USA) in EM buffer with 1mM probenecid (Sigma-Aldrich, St. Louis, MO USA) for 90 or 180 minutes at 4°C or 37°C. Using flow cytometry, live, single cells were gated, and the mean fluorescent intensities of CCF4 fluorescence with 450nm and 535nm emission filters was measured. Ratiometric values of both signals (450 nm : 535 nm, as a ratio of cleaved CCF4 to uncleaved CCF4) was calculated as a means of examining β-lactamase export to the cytosol (Loi et al., 2016).

#### qPCR

*Rubcn^+/+^* and *Rubcn^-/-^* FLT3L-derived-BMDCs were unstimulated or co-cultured with apoptotic Jurkat cells (5 apoptotic cells : 1 DC). RNA was isolated after thirty minutes using the Trizol and RNeasy kit (QIAGEN, Hilden, Germany) according to the manufacturer’s instructions. The RNA was treated with DNase (QIAGEN, Hilden, Germany) to ensure the removal of any contaminating DNA. Extracted RNA was reverse transcribed to cDNA using iScript cDNA synthesis kit (Bio-Rad Laboratories, Hercules, California, U.S.A.). Quantitative, real-time PCR was performed on the QuantStudio 6 Flex Real-Time PCR System (Applied Biosystems, Thermo-Fisher Scientific, Waltham, MA, USA) using TaqMan Universal Master Mix (Applied Biosystems, Waltham, Massachusetts, USA). *Rubcn* (Mm01170688_g1), *Tap 1* (Mm00443188_m1), *Tap 2* (Mm01277033_m1), β*2m* (Mm00437762_m1), and *Actinb* (Mm02619580_g1) TaqMan assay (Life Technologies, Thermo-Fisher Scientific, Waltham, MA, USA) were used. Expression was normalized against *Actinb* to compare target gene mRNA levels.

#### RNA-seq

*Rubcn^+/+^* and *Rubcn^-/-^* FLT3L-derived-BMDCs were unstimulated or co-cultured with apoptotic Jurkat cells (5 apoptotic cells: 1 DC). RNA was isolated after thirty minutes using the Trizol and RNeasy kit (QIAGEN, Hilden, Germany) according to the manufacturer’s instructions. DNase digestion was performed to remove genomic DNA as described above. The sample concentration was measured using the Qubit RNA BR Assay kit (Invitrogen, Carlsbad, CA, USA) and the RNA quality was rechecked with Bioanalyzer 2100 (Agilent, Santa Clara, CA, USA) with RNA Pico chip (Agilent, Santa Clara, CA, USA). 250 ng of RNA was used to make RNA-seq library using Illumina’s TruSeq RNA Library Prep Kit v2. The library concentration was checked using Qubit dsDNA HS assay kit (Invitrogen, Carlsbad, CA, USA) and the library’s fragment size range with Bioanalyzer 2100 using high sensitivity DNA Chip (Agilent, Santa Clara, CA, USA). Then the library was then prepped for sequencing. Sequencing was performed with an Illumina NovaSeq 6000 (Illumina, San Diego, CA, USA) by the NIEHS Epigenomics Core.

Reads were generated as single-end 76mers then filtered to retain only those with an average base quality score of at least 20. Genomic mapping against the mm10 reference genome was performed by STAR v2.5 (Dobin et al., 2013) with the following parameters: --outMultimapperOrder Random --outSAMattrIHstart 0 --outfitter type BySJout --alignSJoverhangMin 8 --limitBAMsortRAM 55000000000 (other parameters at default settings). Mapped read counts per gene were determined by Subread feature Counts v1.5.0-p1 (Liao et al., 2014) with parameter -s0. Differentially expressed genes (DEGs) were identified by DESeq2 v1.14.1 (Love et al., 2014) with filtering threshold FDR 0.05. Gene models for this analysis are based on mm10 RefSeq annotations as downloaded from the UCSC Genome Browser.

The heatmap views for the dataset were generated by heatmap.2 from the gplots R package (R studio, Boston, MA, USA), with row-scaling of the expression values reported by the rlogTransformation function in DESeq2. Ingenuity Pathway Analysis software (Qiagen, Venlo, The Netherlands) was used for further RNA-seq data analysis. The RNA-seq data have been deposited in NCBI’s GEO (https://www.ncbi.nlm.nih.gov/geo/) under accession GSE133945.

#### Western blot

BMDCs were harvested, and the pellets were washed twice with PBS (400 x g, 5 mins, 4 °C). Cells were resuspended in 200 µl of RIPA buffer containing proteinase inhibitor and mixed thoroughly. The cells were lysed on ice for 30 minutes and vortexed for 10 seconds every 10 minutes. For analysis of *Cd11-Cre*-mediated deletion, spleens from *Rubcn^flox/flox^* or *Cd11-Cre^+^ Rubcn^flox/flox^*mice were harvested, and CD11c^+^ populations were isolated using EasySep™ manufacturer’s guidelines from the Mouse Pan-DC Enrichment Kit (STEMCELL Technologies Inc., Vancouver, British Columbia, Canada).

Cell lysates were mixed with 2x Loading buffer, and samples were loaded onto protein gel for separation by molecular weight. Proteins on the gel were transferred onto polyvinylidene difluoride (PVDF) membranes using a Trans-Blot semi-dry transfer cell (Bio-Rad, Hercules, CA USA). The membranes were blocked with Tris-buffered saline containing 0.05% Tween 20 (TBST) and 5% bovine serum albumin (BSA) for 1 hour and incubated with primary antibodies overnight at 4°C. Membranes were washed with PBS containing 0.05% Tween 20 (PBST) or TBST and incubated with horseradish peroxidase (HRP)-conjugated secondary antibodies (Jackson ImmunoResearch, West Grove, PA, USA) for 1 hour, and the HRP on the membranes was developed with ECL Select Western blotting detection reagent and visualized using an ImageQuant LAS 4000 (GE Healthcare, Pittsburgh, PA USA).

#### Flow cytometry

Cells were resuspended in FACs buffer and blocked using anti-mouse CD16/CD32 (Biolegend, San Diego, CA, USA). Zombie UV (Biolegend, San Diego, CA, USA) or Fixable Viability eFluor™ 780 (Thermo-Fisher Scientific, Waltham, MA, USA) dyes were used to stain for cell viability. Cells were stained with FACs antibodies for surface markers, then washed with FACs buffer. 50,000 cells per sample were acquired using the LSRII or Fortessa flow cytometer (BD, Franklin Lakes, NJ, USA) and analyzed using FlowJo software (Tree Star, Ashland, OR, USA). Geometric mean fluorescence intensity (MFI) was calculated for markers of DC activation, such as CD86 and H2-K^b^-OVA_257-264_ (MHC-I SIINFEKL). Antigen-specific T cells were detected using H2-K^b^-OVA_257-264_ Tetramer (NIH Tetramer Core Facility, Atlanta, GA, USA), For Tetramer staining, the samples were incubated with the Tetramer prepared in MACs buffer for 1 hour at room temperature, washed and before additional surface stains.

#### FACs intracellular cytokine staining

For intracellular cytokine staining, cells were incubated with 500 ng/ml PMA (Sigma-Aldrich Corporation, St. Louis, Missouri, United States), 1 µg/ml Ionomycin (Sigma-Aldrich Corporation, St. Louis, Missouri, United States), and 3 µg/ml Brefeldin A (Thermo-Fisher Scientific, Waltham, MA, USA) for 6 hours at 37°C post-surface marker staining. Cells were fixed and permeabilized in BD Cytofix/Cytoperm™ (Becton Dickinson, Franklin Lakes, NJ, USA) per manufacturer’s guidelines, and stained with intracellular antibodies (Perforin, IFNγ, TNFα).

#### Immunofluorescence

BMDCs were plated and stimulated in poly-D-Lysine (1 mg/ml)-coated chamber slides. At indicated time points, cells were fixed with 4% formaldehyde for 20 minutes at 4°C. Following fixation, cells were blocked and permeabilized in block buffer (1% BSA, 0.1% Triton in PBS) for 1 hour at RT. Cells were incubated overnight at 4°C with the primary antibody in block buffer. Cells were washed in TBS-Tween (Tris-buffered saline containing 0.05% Tween-20) and incubated Alexa-Fluor conjugated secondary antibodies (Thermo Fisher Scientific, Waltham, MA, USA). Confocal images were taken with a Zeiss LSM880 (Carl Zeiss Inc, Oberkochen, Germany) using the 488nm, 561nm, and 633nm laser lines for excitation paired with emission filter setting of 495-558nm, 571-624nm, and 640-758nm respectively. An EC Plan-Neofluar 40X/0.9 M27 objective was used for image collection along with a pinhole setting of 3.4 Airy Units which yields an optical slice of 3.2um.

#### Quantification of immunofluorescence

ImageJ Fiji software (Schindelin et al., 2012) was used to quantitate the fluorescence intensity across the bead-containing phagosome. The images were imported into the Fiji software and a region of interest was selected to determine the mean fluorescent intensity. Data was compiled from 10-15 cells per treatment over three independent experiments.

#### Live cell imaging

1 x10^6^ DCs were seeded on 50 µg/ml poly-D-Lysine ((Sigma-Aldrich Corporation, St. Louis, Missouri, USA)-precoated 35 mm, Petri dish (MatTek Corporation, Ashland, MA). 5 x10^6^ PKH26-labeled apoptotic B16-OVA cells were added to the culture. The cells were maintained at ∼37 ◦C and 5% CO2 using an environmental control chamber. Confocal images were taken with a Zeiss LSM880 (Carl Zeiss Inc, Oberkochen, Germany) using the 561nm laser line for excitation paired with emission filter settings of 571-624nm. A Plan-APOCHROMAT 63X/1.4 Oil DIC objective was used for image collection along with a pinhole setting of 2.3 Airy units which yields an optical slice of 2 µm.

#### Quantification of cytokine and chemokine production

Supernatants from DC cultures were collected 18 hours after co-culture with apoptotic B16-OVA cells (5 apoptotic cells : 1 DC). The Bio-Plex® Multiplex Immunoassay System (Bio-Rad, Hercules, CA) was used following the manufacturer’s instructions to detect the cytokines and chemokines released from the co-cultures. IFNβ was detected using Legend Max IFNβ ELISA kit (Biolegend, San Diego, CA, US). Data (mean ± SEM) represent three independent experiments from 5 mice per genotype.

#### Statistical Analysis

All data were statistically analyzed using GraphPad Prism software v6.01 (GraphPad Software, La Jolla, CA, USA). The statistical significance was determined by a Student t-test or two-way ANOVA with multiple comparison test (MCT) as indicated in the figure legends and p< 0.05 was considered significant.

## Supplemental Information Legends

**Figure S1, related to Figure 1:**
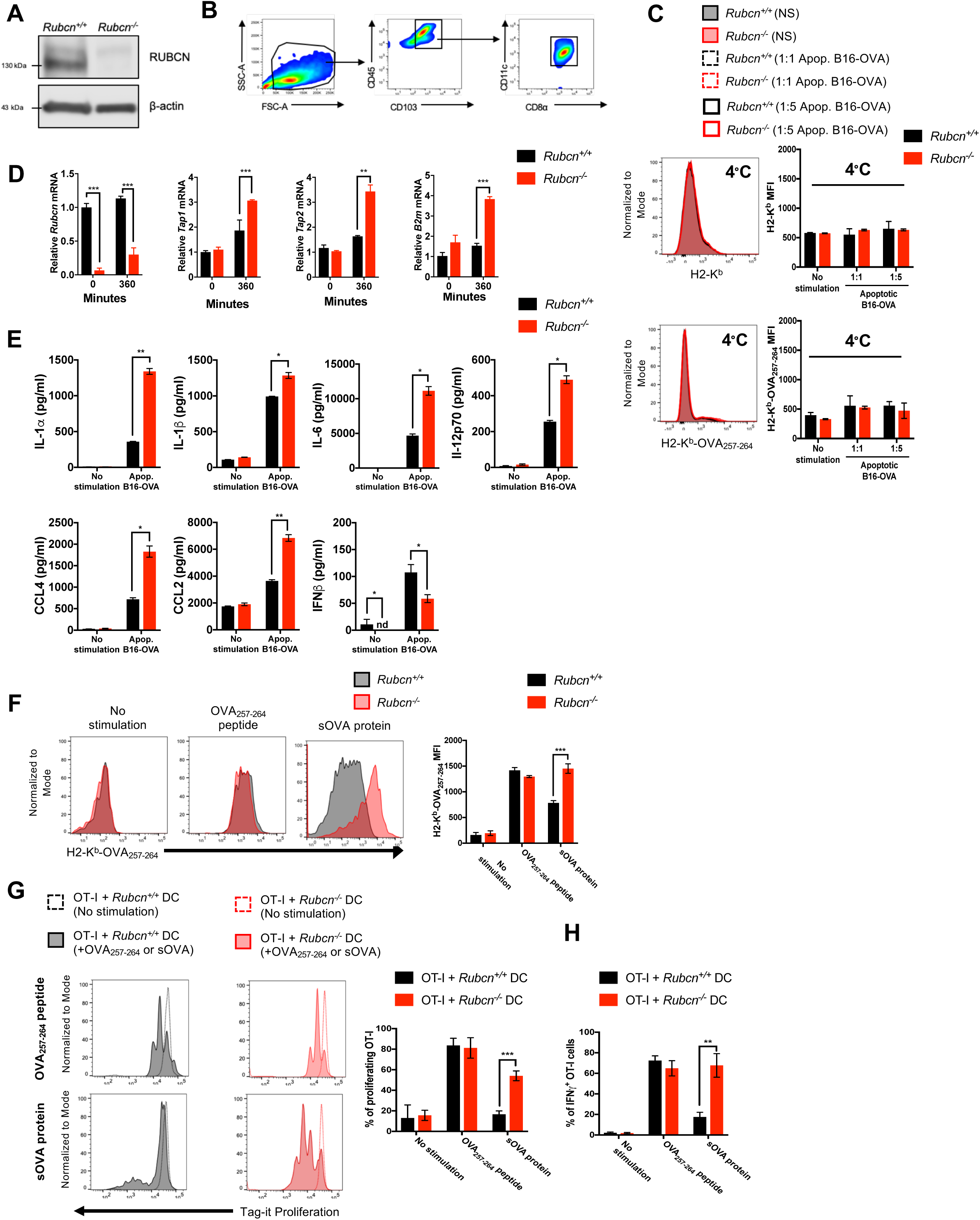
*Rubcn^-/-^* DCs display increased cross-presentation capacity but equivalent levels of peptide-mediated MHCI antigen presentation. Bone marrow-derived dendritic cells (DCs) were generated from *Rubcn^+/+^*(black) and *Rubcn^-/-^* (red) mice *in vitro* with FLT3-L for 7 days. (**A**) DCs were harvested for Western blot analysis of RUBCN and β-actin expression. (**B**) Representative flow cytometry plots for DCs on day 7 post-differentiation. (**C**) DCs were co-cultured with apoptotic B16-OVA cells (1 or 5 apoptotic cells: 1 DC) at 4°C. Eighteen hours later, DCs were harvested for flow cytometry analysis of H2-K^b^ and H2-K^b^-OVA_257-264_ expression. (**D**) DCs were co-cultured with apoptotic Jurkat cells (5 apoptotic cells: 1 DC). Six hours later, RNA was harvested for qPCR of cross-presentation machinery, *Tap1*, *Tap2*, *B2m*, and *Rubcn*. Data are normalized to *Actinb* expression and expressed as mean ± SEM. (**E**) DCs were co-cultured with apoptotic B16-OVA cells (5 apoptotic cells: 1 DC). Eighteen hours later, co-culture supernatant was collected and analyzed for cytokine and chemokine production by multiplex or ELISA technology. (**F**) DCs were co-cultured with OVA_257-264_ peptide (50 ng/ml) or sOVA protein (150 μg/ml). Eighteen hours later, DCs were harvested for flow cytometry analysis of H2-K^b^-OVA_257-264_ expression. (**G-H**) DCs were co-cultured with OVA_257-264_ peptide (50 ng/ml) or sOVA protein (150 μg/ml). Eighteen hours later, loaded DCs were co-cultured with naïve OT-I CD8^+^ T cells, labeled with Tag-it Proliferation dye. Forty-eight hours later, *in vitro* cultures were harvested for flow cytometry analysis of OT-I proliferation (**G**) or IFNγ production (**H**). Data are expressed as mean ± SEM. No less than three independent experiments were performed, with 3-5 replicates per condition. Significance was calculated using 2-way ANOVA (*p<0.05, **p<0.01, ***p<0.001).

**Figure S2, related to Figure 1:**
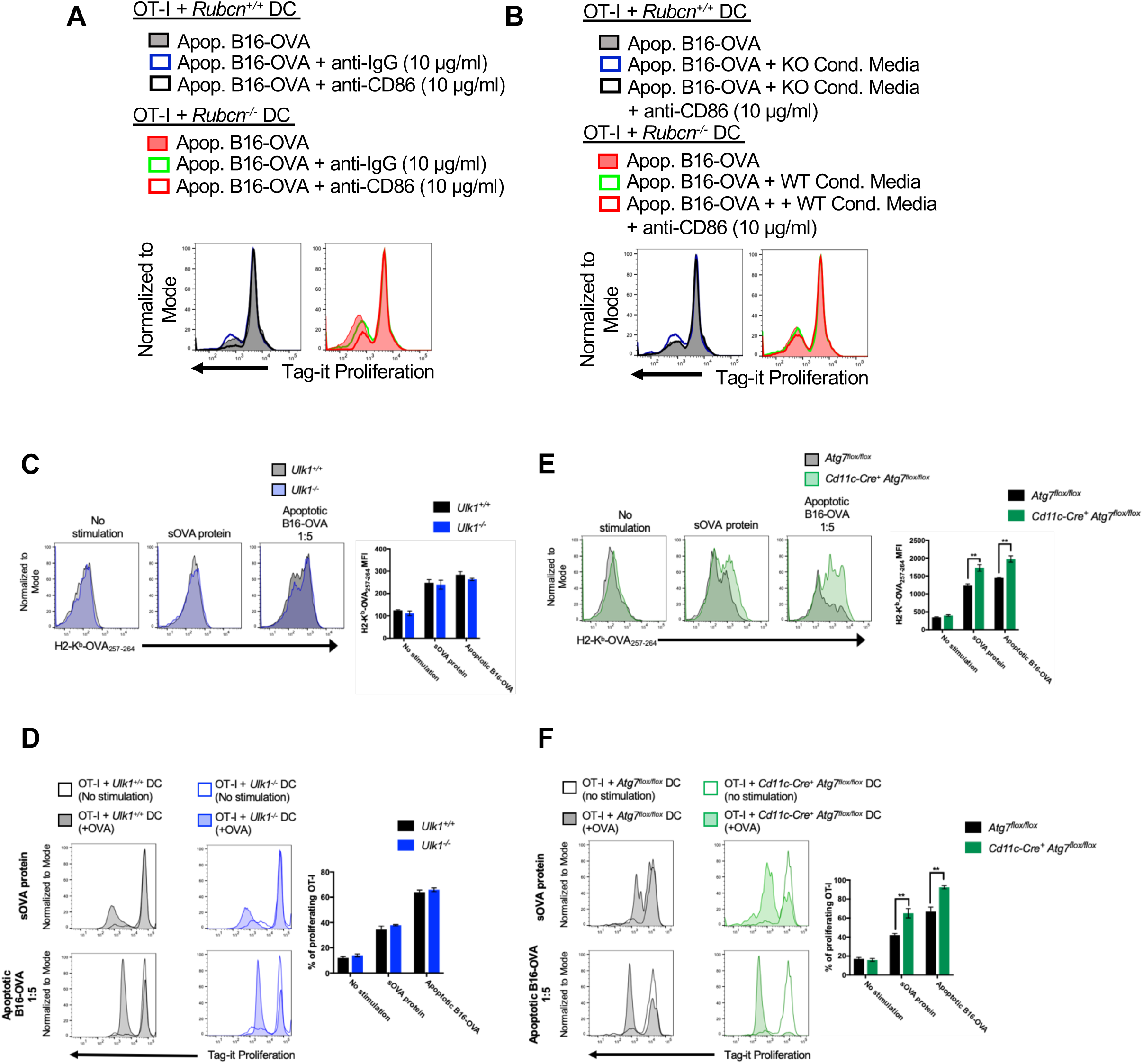
Defects in LAP, not canonical autophagy, results in increased cross-presentation. (**A-B**) Bone marrow-derived dendritic cells (DCs) were generated from *Rubcn^+/+^* (black) and *Rubcn^-/-^* (red) mice *in vitro* with FLT3-L for 7 days. DCs were co-cultured with apoptotic B16-OVA cells (5 apoptotic cells: 1 DC). Eighteen hours later, loaded DCs incubated with anti-IgG, anti-CD86, and/or conditioned media from independent cultures, as indicated for at least 1 hour. DCs were then were co-cultured with naïve OT-I CD8^+^ T cells, labeled with Tag-it Proliferation dye. Forty-eight hours later, *in vitro* cultures were harvested for flow cytometry flow cytometry analysis of OT-I proliferation. Representative histograms are shown. (**C-D**). Bone marrow-derived dendritic cells (DCs) were generated from *Ulk1^+/+^* (black) and *Ulk1^-/-^* (blue) mice *in vitro* with FLT3-L for 7 days. DCs were co-cultured with sOVA protein (150 μg/ml) or apoptotic B16-OVA cells (5 apoptotic cells: 1 DC). Eighteen hours later, DCs were harvested for flow cytometry analysis of H2-K^b^-OVA_257-264_ expression (**C**). DCs were co-cultured sOVA protein (150 μg/ml) or apoptotic B16-OVA cells (5 apoptotic cells: 1 DC). Eighteen hours later, loaded DCs were co-cultured with naïve OT-I CD8^+^ T cells, labeled with Tag-it Proliferation dye. Forty-eight hours later, *in vitro* cultures were harvested for flow cytometry analysis of OT-I proliferation (**D**). (**E-F**). Bone marrow-derived dendritic cells (DCs) were generated from *Atg7^flox/flox^* (black) and *Cd11c-Cre^+^ Atg7^flox/flox^* (green) mice *in vitro* with FLT3-L for 7 days. DCs were co-cultured with sOVA protein (150 μg/ml) or apoptotic B16-OVA cells (5 apoptotic cells: 1 DC). Eighteen hours later, DCs were harvested for flow cytometry analysis of H2-K^b^-OVA_257-264_ expression (**E**). DCs were co-cultured sOVA protein (150 μg/ml) or apoptotic B16-OVA cells (5 apoptotic cells: 1 DC). Eighteen hours later, loaded DCs were co-cultured with naïve OT-I CD8^+^ T cells, labeled with Tag-it Proliferation dye. Forty-eight hours later, *in vitro* cultures were harvested for flow cytometry analysis of OT-I proliferation (**F**). Data are expressed as mean ± SEM. No less than three independent experiments were performed, with 3-5 replicates per condition. Significance was calculated using 2-way ANOVA (*p<0.05, **p<0.01, ***p<0.001).

**Figure S3, related to Figures 2 and 3:**
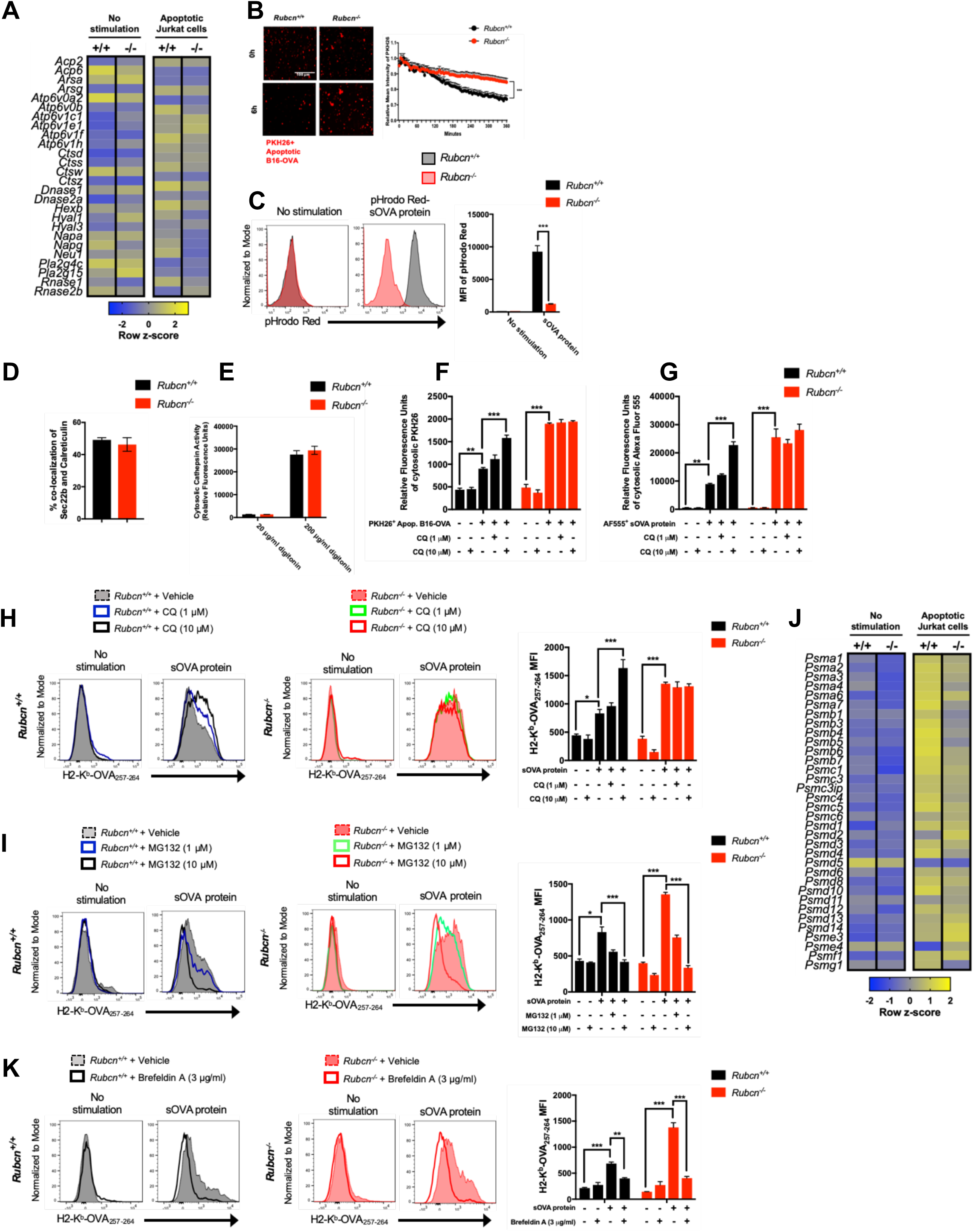
*Rubcn^-/-^* DCs display increased phagosome-to-cytosol escape and sensitive to proteasome inhibition. Bone marrow-derived dendritic cells (DCs) were generated from *Rubcn^+/+^*(black) and *Rubcn^-/-^* (red) mice *in vitro* with FLT3-L for 7 days. (**A**) DCs were co-cultured with apoptotic Jurkat cells (5 apoptotic cells: 1 DC). Thirty minutes later, RNA was harvested for RNA-seq. Genes with fold change (FC>1.25) compared to no stimulation and p<0.05 (ANOVA) were considered significant and entered in Ingenuity Pathway Analysis (IPA) and R studio for further pathway analysis. Heatmap of lysosomal machinery depicted as average row z-score is shown (n=3 per genotype). (**B**) DCs were co-cultured with PKH26-labeled apoptotic B16-OVA cells (5 apoptotic cells: 1 DC). Time-lapse imaging was performed for 6 hours to examine the rate of clearance of apoptotic cells. Mean fluorescence of PKH26 per field was obtained from images using Fiji. Data are expressed as mean ± SEM. (n=15 fields per timepoint per condition). (**C**) sOVA protein was labeled with pHrodo Red per manufacturer’s instructions, then co-cultured (150 µg/ml) with *Rubcn^+/+^* and *Rubcn^-/-^*DCs *in vitro*. Four hours later, pHrodo Red fluorescence was measured in DCs by flow cytometry, as a readout of phagosomal maturation. (**D**) DCs were plated in in chamber slides, and were fixed, permeabilized, stained for Sec22b (blue), H2-K^b^ (red), and Calreticulin (green), and imaged via confocal microscopy. Co-localization of Sec22b and Calreticulin was obtained from images using Fiji. Data are expressed as mean ± SEM (n=15 cells per condition). (**E**) DCs were permeabilized with 20 µg/ml or 200 µg/ml digitonin, then incubated with cathepsin-specific fluorogenic substrate, Z-FR-AMC (30 µM), which emits fluorescence at 460 nm upon cathepsin-mediated cleavage. Membrane portions of samples were pelleted by centrifugation, and cytosolic supernatant portions were collected and analyzed for fluorescence (460 nm) using Tecan microplate reader. (**F-G**). DCs were co-cultured with PKH26-labeled apoptotic B16-OVA cells (5 apoptotic cells: 1 DC, **F**) or Alexa Fluor 555-labeled sOVA protein (150 µg/ml, **G**). Six hours later, DCs were permeabilized with digitonin (20 µg/ml), and membrane portions of samples were pelleted by centrifugation. Cytosolic supernatant portions were collected and analyzed for fluorescence (567 nm [**F**] or 580 nm [**G**]) using Tecan microplate reader. (**H**) DCs were pre-treated with vehicle or chloroquine (CQ) at 1 or 10 µM for 2 hours and then co-cultured with sOVA protein (150 µg/ml). Eighteen hours later, DCs were harvested for flow cytometry analysis of H2-K^b^-OVA_257-264_ expression. (**I**) DCs were pre-treated with vehicle or MG-132 at 1 or 10 µM for 2 hours and then co-cultured in fresh media with sOVA protein (150 µg/ml). Eighteen hours later, DCs were harvested for flow cytometry analysis of H2-K^b^-OVA_257-264_ expression. (**J**) DCs were co-cultured with apoptotic Jurkat cells (5 apoptotic cells: 1 DC). Thirty minutes later, RNA was harvested for RNA-seq. Genes with fold change (FC>1.25) compared to no stimulation and p<0.05 (ANOVA) were considered significant and entered in Ingenuity Pathway Analysis (IPA) and R studio for further pathway analysis. Heatmap of proteasome machinery depicted as average row z-score is shown (n=3 per genotype). (**K**) DCs were pre-treated with vehicle or Brefeldin A at 3 µg/ml for 2 hours and then co-cultured in fresh media with sOVA protein (150 µg/ml). Eighteen hours later, DCs were harvested for flow cytometry analysis of H2-K^b^-OVA_257-264_ expression. Data are expressed as mean ± SEM. No less than two independent experiments were performed, with 3-5 replicates per condition. Significance was calculated using 2-way ANOVA (*p<0.05, **p<0.01, ***p<0.001).

**Figure S4, related to Figure 4:**
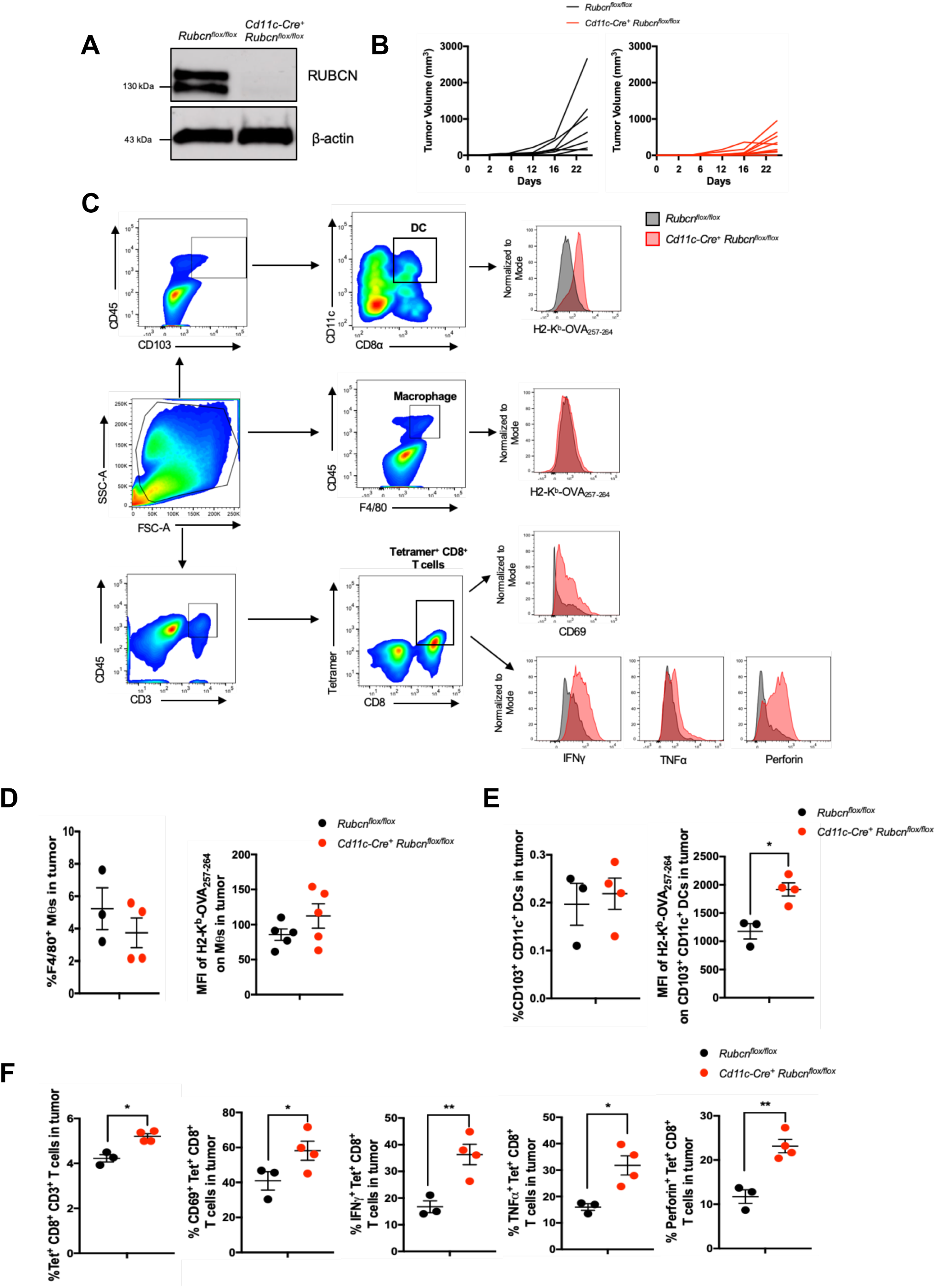
*Cd11c-Cre Rubcn^flox/^*^flox^ mice display increased anti-tumor immunity at day 22 *in vivo.* (**A**) Spleens from *Rubcn^flox/flox^*and *Cd11-Cre^+^ Rubcn^flox/flox^* littermates were harvested. CD11c^+^ cells were isolated via magnetic bead separation and subjected to Western blot analysis for RUBCN and β-actin expression. (**B**) 1 x 10^6^ B16-OVA cells were injected intradermally into *Rubcn^flox/flox^*(black, n=7) or *Cd11-Cre^+^ Rubcn^flox/flox^* (red, n=8) mice. Tumor growth for individual mice was monitored and recorded. (**C**) 1 x 10^6^ B16-OVA cells were injected intradermally into *Rubcn^flox/flox^* (black, n=5) or *Cd11-Cre^+^ Rubcn^flox/flox^* (red, n=5) mice. On day 16 post-injection, tumors harvested for flow cytometry analysis. Representative flow cytometry plots and gating strategies for tumor-infiltrating immune cells are shown. **(D**) On day 16 post-injection, tumors were harvested for flow cytometry analysis. Percentage of F480^+^ macrophages (M*θ*s) within CD45^+^ population and MFI of H2-K^b^-OVA_257-264_ expression are expressed as mean ± SEM. (**E-F**) On day 22 post-injection, tumors were harvested for flow cytometry analysis.. Percentage of CD11c^+^ CD103^+^ CD8α^+^ DCs within CD45^+^ population and MFI of H2-K^b^-OVA_257-264_ expression (**E**) are expressed as mean ± SEM. Percentage of CD8α^+^ Tetramer^+^ T cells within CD45^+^ population and percentage of CD69^+^, IFN*γ*^+^, TNFα^+^, and Perforin^+^ CD8+ T cells within tumor (**F**) are expressed as mean ± SEM. No less than three independent experiments were performed, with 5 replicates per genotype. Significance was calculated using 2-way ANOVA (*p<0.05, **p<0.01, ***p<0.001).

## STAR METHODS

### KEY RESOURCES TABLE

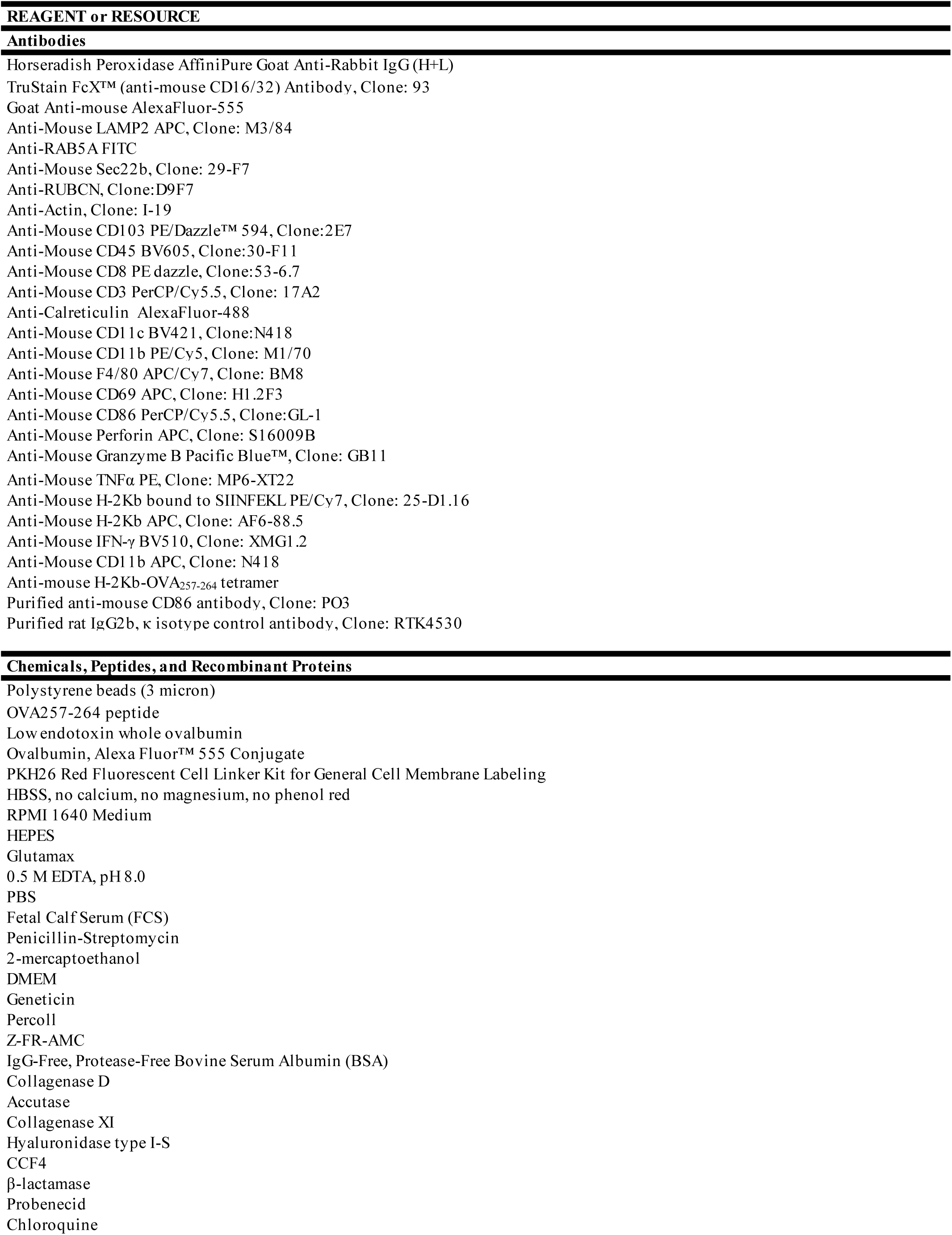
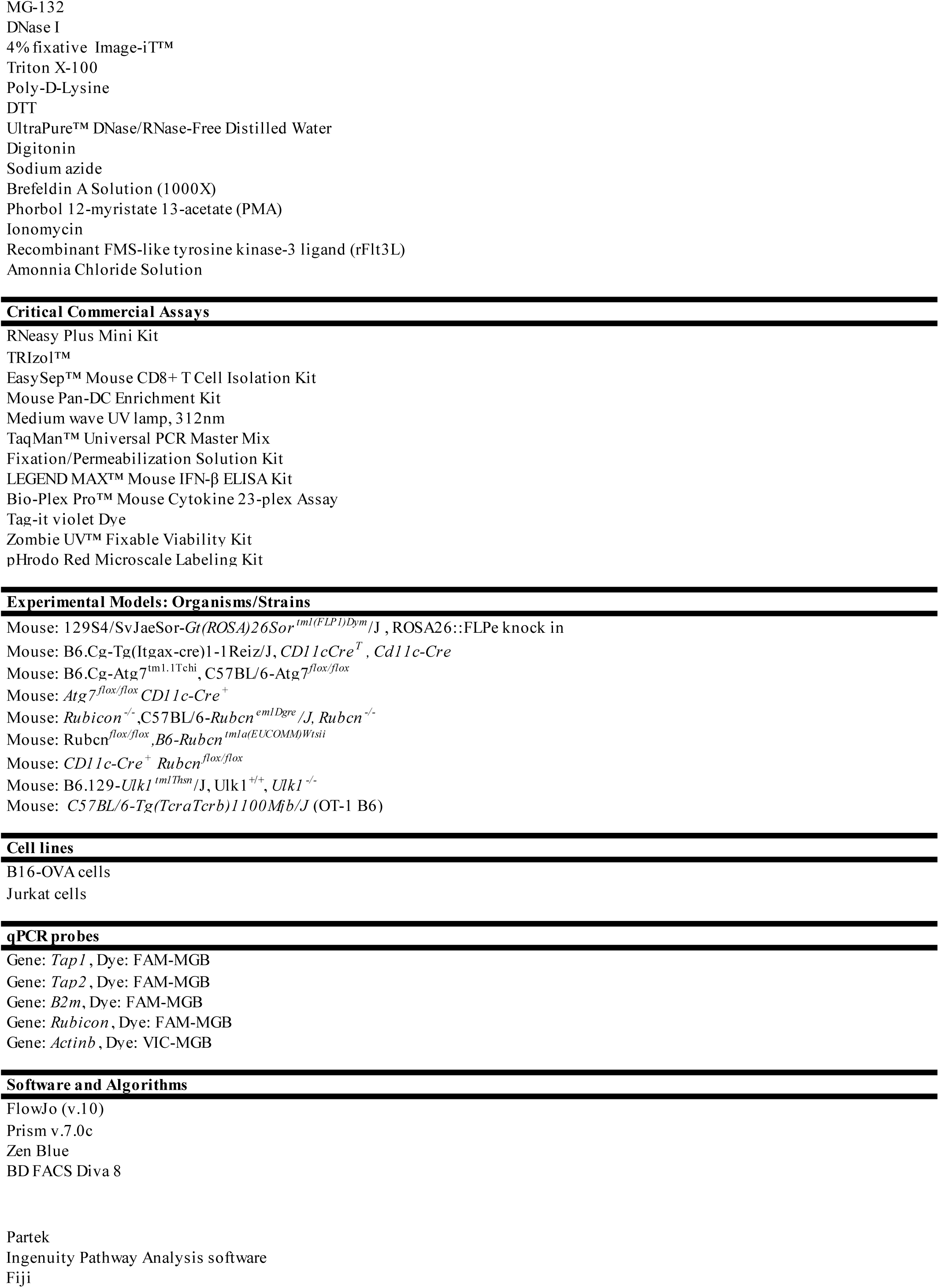

---

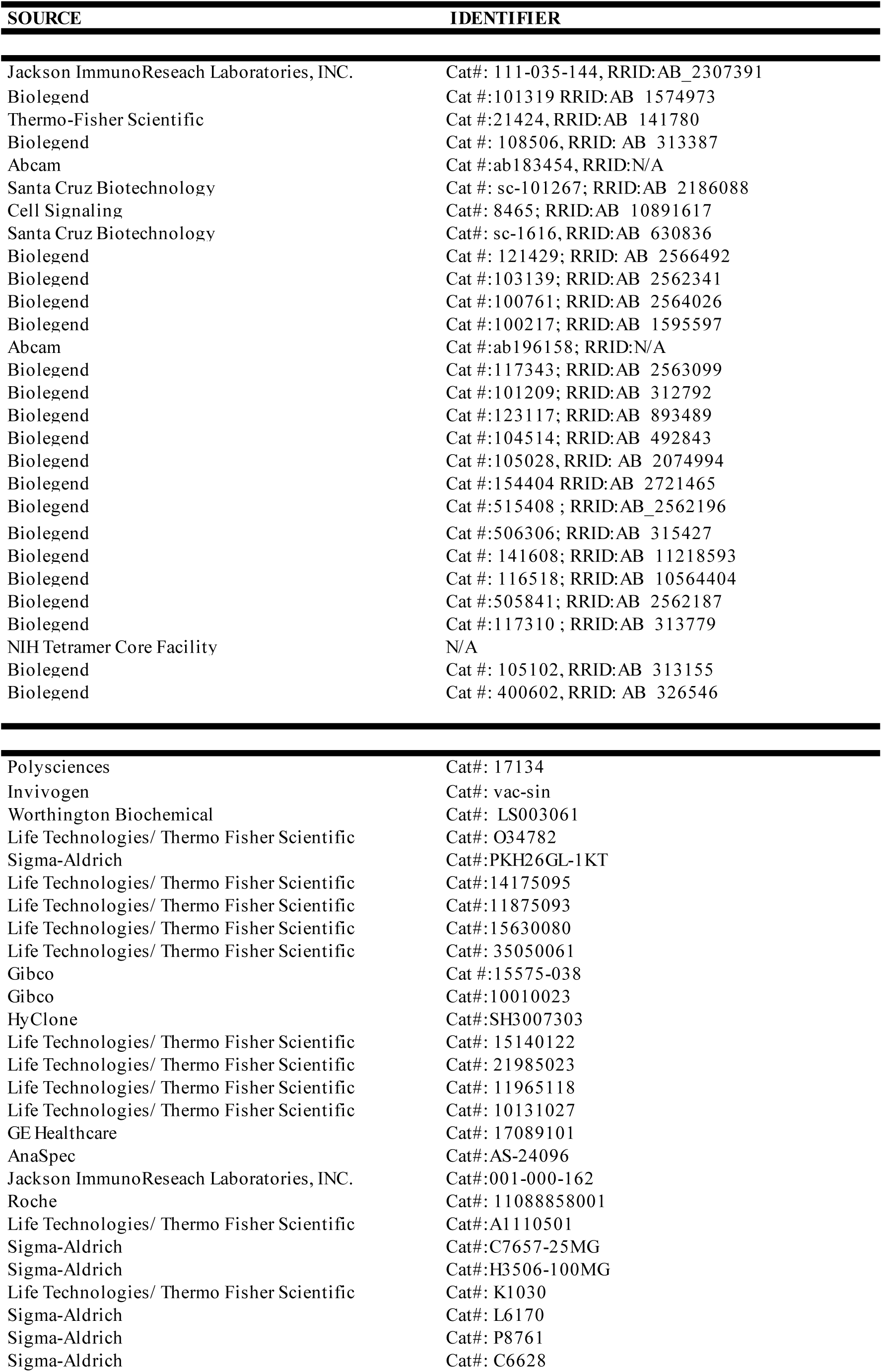
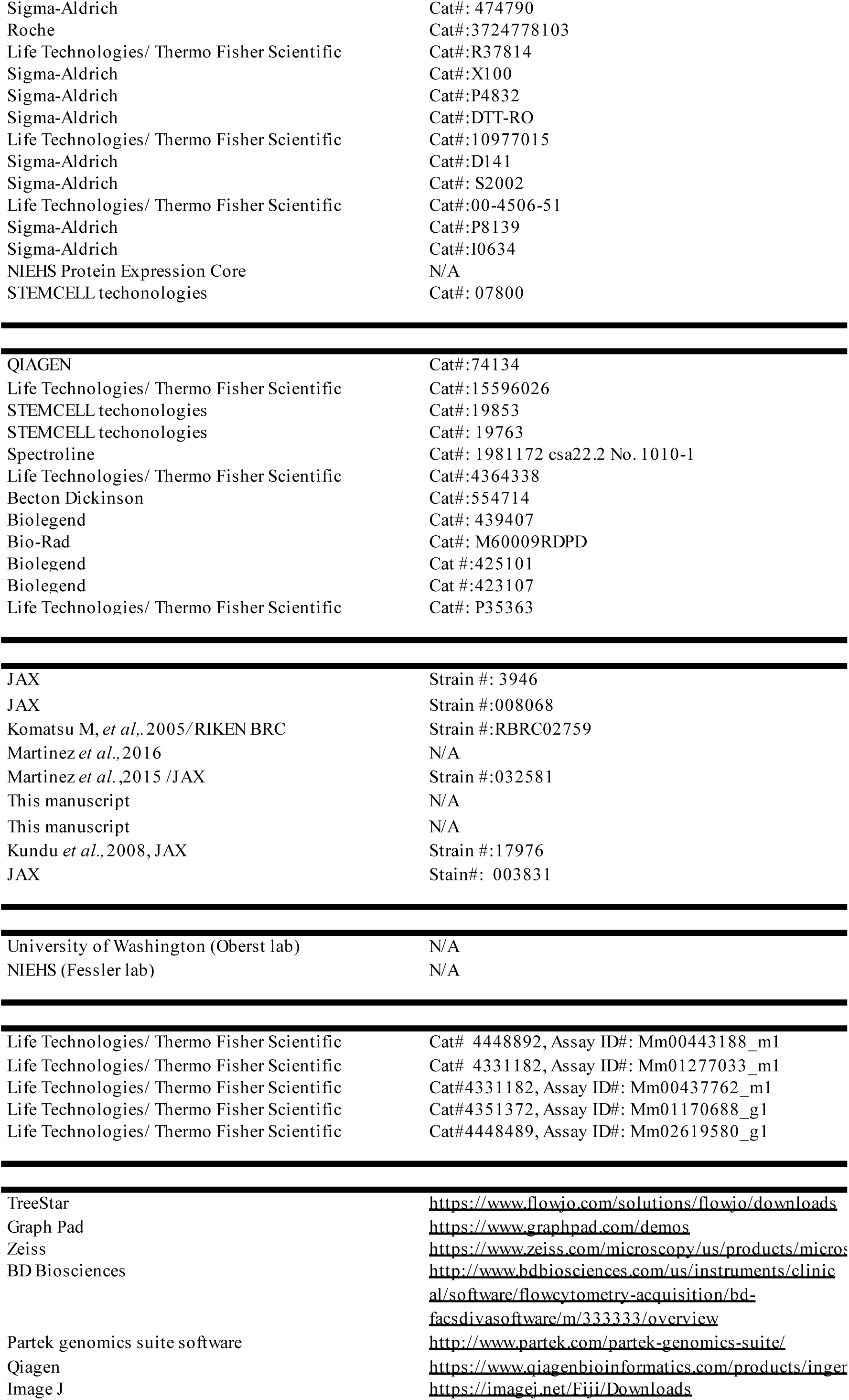

2005. 169(3): p. 425-34.

maturation. Blood, 2008. 112(4): p. 1493-502.

and autophagy proteins. Nat Cell Biol, 2015. 17(7): p. 893-906.

016. 533(7601): p. 115-9.

products/microscope-software/zen.html

products/ingenuity-pathway-analysis/

## References

Alloatti, A., Rookhuizen, D.C., Joannas, L., Carpier, J.M., Iborra, S., Magalhaes, J.G., Yatim, N., Kozik, P., Sancho, D., Albert, M.L., and Amigorena, S. (2017). Critical role for Sec22b-dependent antigen cross-presentation in antitumor immunity. J Exp Med 214, 2231–2241.

Andrieu, M., Desoutter, J.F., Loing, E., Gaston, J., Hanau, D., Guillet, J.G., and Hosmalin, A. (2003). Two human immunodeficiency virus vaccinal lipopeptides follow different cross-presentation pathways in human dendritic cells. J Virol 77, 1564–1570.

Belizaire, R., and Unanue, E.R. (2009). Targeting proteins to distinct subcellular compartments reveals unique requirements for MHC class I and II presentation. Proc Natl Acad Sci U S A 106, 17463–17468.

Blachere, N.E., Darnell, R.B., and Albert, M.L. (2005). Apoptotic cells deliver processed antigen to dendritic cells for cross-presentation. PLoS Biol 3, e185.

Boya, P., Gonzalez-Polo, R.A., Poncet, D., Andreau, K., Vieira, H.L., Roumier, T., Perfettini, J.L., and Kroemer, G. (2003). Mitochondrial membrane permeabilization is a critical step of lysosome-initiated apoptosis induced by hydroxychloroquine. Oncogene 22, 3927–3936.

Brasel, K., De Smedt, T., Smith, J.L., and Maliszewski, C.R. (2000). Generation of murine dendritic cells from flt3-ligand-supplemented bone marrow cultures. Blood 96, 3029–3039.

Burgdorf, S., Lukacs-Kornek, V., and Kurts, C. (2006). The mannose receptor mediates uptake of soluble but not of cell-associated antigen for cross-presentation. J Immunol 176, 6770–6776.

Burgdorf, S., Scholz, C., Kautz, A., Tampe, R., and Kurts, C. (2008). Spatial and mechanistic separation of cross-presentation and endogenous antigen presentation. Nat Immunol 9, 558–566.

Calderon, B., and Unanue, E.R. (2012). Antigen presentation events in autoimmune diabetes. Curr Opin Immunol 24, 119–128.

Cebrian, I., Visentin, G., Blanchard, N., Jouve, M., Bobard, A., Moita, C., Enninga, J., Moita, L.F., Amigorena, S., and Savina, A. (2011). Sec22b regulates phagosomal maturation and antigen crosspresentation by dendritic cells. Cell 147, 1355–1368.

Cruz, F.M., Colbert, J.D., Merino, E., Kriegsman, B.A., and Rock, K.L. (2017). The Biology and Underlying Mechanisms of Cross-Presentation of Exogenous Antigens on MHC-I Molecules. Annu Rev Immunol 35, 149–176.

Cunha, L.D., Yang, M., Carter, R., Guy, C., Harris, L., Crawford, J.C., Quarato, G., Boada-Romero, E., Kalkavan, H., Johnson, M.D.L., et al. (2018). LC3-Associated Phagocytosis in Myeloid Cells Promotes Tumor Immune Tolerance. Cell 175, 429–441 e416.

Dobin, A., Davis, C.A., Schlesinger, F., Drenkow, J., Zaleski, C., Jha, S., Batut, P., Chaisson, M., and Gingeras, T.R. (2013). STAR: ultrafast universal RNA-seq aligner. Bioinformatics 29, 15–21.

Fehres, C.M., Unger, W.W., Garcia-Vallejo, J.J., and van Kooyk, Y. (2014). Understanding the biology of antigen cross-presentation for the design of vaccines against cancer. Front Immunol 5, 149.

Garcia-Mata, R., Szul, T., Alvarez, C., and Sztul, E. (2003). ADP-ribosylation factor/COPI-dependent events at the endoplasmic reticulum-Golgi interface are regulated by the guanine nucleotide exchange factor GBF1. Mol Biol Cell 14, 2250–2261.

Greenwald, R.J., Freeman, G.J., and Sharpe, A.H. (2005). The B7 family revisited. Annu Rev Immunol 23, 515–548.

Gutierrez-Martinez, E., Planes, R., Anselmi, G., Reynolds, M., Menezes, S., Adiko, A.C., Saveanu, L., and Guermonprez, P. (2015). Cross-Presentation of Cell-Associated Antigens by MHC Class I in Dendritic Cell Subsets. Front Immunol 6, 363.

Hansen, T.H., and Bouvier, M. (2009). MHC class I antigen presentation: learning from viral evasion strategies. Nat Rev Immunol 9, 503–513.

Hayashi, K., Taura, M., and Iwasaki, A. (2018). The interaction between IKKalpha and LC3 promotes type I interferon production through the TLR9-containing LAPosome. Sci Signal 11.

Heckmann, B.L., Teubner, B.J.W., Tummers, B., Boada-Romero, E., Harris, L., Yang, M., Guy, C.S., Zakharenko, S.S., and Green, D.R. (2019). LC3-Associated Endocytosis Facilitates beta-Amyloid Clearance and Mitigates Neurodegeneration in Murine Alzheimer’s Disease. Cell 178, 536–551 e514.

Henault, J., Martinez, J., Riggs, J.M., Tian, J., Mehta, P., Clarke, L., Sasai, M., Latz, E., Brinkmann, M.M., Iwasaki, A., et al. (2012). Noncanonical autophagy is required for type I interferon secretion in response to DNA-immune complexes. Immunity 37, 986–997.

Hogquist, K.A., Jameson, S.C., Heath, W.R., Howard, J.L., Bevan, M.J., and Carbone, F.R. (1994). T cell receptor antagonist peptides induce positive selection. Cell 76, 17–27.

Huang, A.Y., Golumbek, P., Ahmadzadeh, M., Jaffee, E., Pardoll, D., and Levitsky, H. (1994). Role of bone marrow-derived cells in presenting MHC class I-restricted tumor antigens. Science 264, 961–965.

Jantas, D., Greda, A., Leskiewicz, M., Grygier, B., Pilc, A., and Lason, W. (2015). Neuroprotective effects of mGluR II and III activators against staurosporine- and doxorubicin-induced cellular injury in SH-SY5Y cells: New evidence for a mechanism involving inhibition of AIF translocation. Neurochem Int 88, 124–137.

Ji, Q., Castelli, L., and Goverman, J.M. (2013). MHC class I-restricted myelin epitopes are cross-presented by Tip-DCs that promote determinant spreading to CD8(+) T cells. Nat Immunol 14, 254–261.

Joffre, O.P., Segura, E., Savina, A., and Amigorena, S. (2012). Cross-presentation by dendritic cells. Nat Rev Immunol 12, 557–569.

Kantoff, P.W., Higano, C.S., Shore, N.D., Berger, E.R., Small, E.J., Penson, D.F., Redfern, C.H., Ferrari, A.C., Dreicer, R., Sims, R.B., et al. (2010). Sipuleucel-T immunotherapy for castration-resistant prostate cancer. N Engl J Med 363, 411–422.

Kaufman, J. (2018). Generalists and Specialists: A New View of How MHC Class I Molecules Fight Infectious Pathogens. Trends Immunol 39, 367–379.

Keller, C., Mellouk, N., Danckaert, A., Simeone, R., Brosch, R., Enninga, J., and Bobard, A. (2013). Single cell measurements of vacuolar rupture caused by intracellular pathogens. J Vis Exp, e50116.

Keller, C.W., Sina, C., Kotur, M.B., Ramelli, G., Mundt, S., Quast, I., Ligeon, L.A., Weber, P., Becher, B., Munz, C., and Lunemann, J.D. (2017). ATG-dependent phagocytosis in dendritic cells drives myelin-specific CD4(+) T cell pathogenicity during CNS inflammation. Proc Natl Acad Sci U S A 114, E11228–E11237.

Komatsu, M., Waguri, S., Ueno, T., Iwata, J., Murata, S., Tanida, I., Ezaki, J., Mizushima, N., Ohsumi, Y., Uchiyama, Y., et al. (2005). Impairment of starvation-induced and constitutive autophagy in Atg7-deficient mice. J Cell Biol 169, 425–434.

Kovacsovics-Bankowski, M., and Rock, K.L. (1995). A phagosome-to-cytosol pathway for exogenous antigens presented on MHC class I molecules. Science 267, 243–246.

Kundu, M., Lindsten, T., Yang, C.Y., Wu, J., Zhao, F., Zhang, J., Selak, M.A., Ney, P.A., and Thompson, C.B. (2008). Ulk1 plays a critical role in the autophagic clearance of mitochondria and ribosomes during reticulocyte maturation. Blood 112, 1493–1502.

Liao, Y., Smyth, G.K., and Shi, W. (2014). featureCounts: an efficient general purpose program for assigning sequence reads to genomic features. Bioinformatics 30, 923–930.

Loi, M., Muller, A., Steinbach, K., Niven, J., Barreira da Silva, R., Paul, P., Ligeon, L.A., Caruso, A., Albrecht, R.A., Becker, A.C., et al. (2016). Macroautophagy Proteins Control MHC Class I Levels on Dendritic Cells and Shape Anti-viral CD8(+) T Cell Responses. Cell Rep 15, 1076–1087.

Love, M.I., Huber, W., and Anders, S. (2014). Moderated estimation of fold change and dispersion for RNA-seq data with DESeq2. Genome biology 15, 550.

Maranon, C., Desoutter, J.F., Hoeffel, G., Cohen, W., Hanau, D., and Hosmalin, A. (2004). Dendritic cells cross-present HIV antigens from live as well as apoptotic infected CD4+ T lymphocytes. Proc Natl Acad Sci U S A 101, 6092–6097.

Marino, E., Tan, B., Binge, L., Mackay, C.R., and Grey, S.T. (2012). B-cell cross-presentation of autologous antigen precipitates diabetes. Diabetes 61, 2893–2905.

Martinez, J., Almendinger, J., Oberst, A., Ness, R., Dillon, C.P., Fitzgerald, P., Hengartner, M.O., and Green, D.R. (2011a). Microtubule-associated protein 1 light chain 3 alpha (LC3)-associated phagocytosis is required for the efficient clearance of dead cells. Proceedings of the National Academy of Sciences of the United States of America 108, 17396–17401.

Martinez, J., Almendinger, J., Oberst, A., Ness, R., Dillon, C.P., Fitzgerald, P., Hengartner, M.O., and Green, D.R. (2011b). Microtubule-associated protein 1 light chain 3 alpha (LC3)-associated phagocytosis is required for the efficient clearance of dead cells. Proc Natl Acad Sci U S A 108, 17396–17401.

Martinez, J., Cunha, L.D., Park, S., Yang, M., Lu, Q., Orchard, R., Li, Q.Z., Yan, M., Janke, L., Guy, C., et al. (2016a). Corrigendum: Noncanonical autophagy inhibits the autoinflammatory, lupus-like response to dying cells. Nature 539, 124.

Martinez, J., Cunha, L.D., Park, S., Yang, M., Lu, Q., Orchard, R., Li, Q.Z., Yan, M., Janke, L., Guy, C., et al. (2016b). Noncanonical autophagy inhibits the autoinflammatory, lupus-like response to dying cells. Nature 533, 115–119.

Martinez, J., Malireddi, R.K., Lu, Q., Cunha, L.D., Pelletier, S., Gingras, S., Orchard, R., Guan, J.L., Tan, H., Peng, J., et al. (2015). Molecular characterization of LC3-associated phagocytosis reveals distinct roles for Rubicon, NOX2 and autophagy proteins. Nat Cell Biol 17, 893–906.

Mauthe, M., Orhon, I., Rocchi, C., Zhou, X., Luhr, M., Hijlkema, K.J., Coppes, R.P., Engedal, N., Mari, M., and Reggiori, F. (2018). Chloroquine inhibits autophagic flux by decreasing autophagosome-lysosome fusion. Autophagy 14, 1435–1455.

Nair-Gupta, P., Baccarini, A., Tung, N., Seyffer, F., Florey, O., Huang, Y., Banerjee, M., Overholtzer, M., Roche, P.A., Tampe, R., et al. (2014). TLR signals induce phagosomal MHC-I delivery from the endosomal recycling compartment to allow cross-presentation. Cell 158, 506–521.

Palmowski, M.J., Gileadi, U., Salio, M., Gallimore, A., Millrain, M., James, E., Addey, C., Scott, D., Dyson, J., Simpson, E., and Cerundolo, V. (2006). Role of immunoproteasomes in cross-presentation. J Immunol 177, 983–990.

Pooley, J.L., Heath, W.R., and Shortman, K. (2001). Cutting edge: intravenous soluble antigen is presented to CD4 T cells by CD8-dendritic cells, but cross-presented to CD8 T cells by CD8+ dendritic cells. J Immunol 166, 5327–5330.

Rahimi, R.A., and Luster, A.D. (2018). Chemokines: Critical Regulators of Memory T Cell Development, Maintenance, and Function. Adv Immunol 138, 71–98.

Repnik, U., Cesen, M.H., and Turk, B. (2016). Measuring Cysteine Cathepsin Activity to Detect Lysosomal Membrane Permeabilization. Cold Spring Harb Protoc 2016.

Rock, K.L., Reits, E., and Neefjes, J. (2016). Present Yourself! By MHC Class I and MHC Class II Molecules. Trends Immunol 37, 724–737.

Rouillard, A.D., Gundersen, G.W., Fernandez, N.F., Wang, Z., Monteiro, C.D., McDermott, M.G., and Ma’ayan, A. (2016). The harmonizome: a collection of processed datasets gathered to serve and mine knowledge about genes and proteins. Database (Oxford) 2016.

Sanchez-Paulete, A.R., Cueto, F.J., Martinez-Lopez, M., Labiano, S., Morales-Kastresana, A., Rodriguez-Ruiz, M.E., Jure-Kunkel, M., Azpilikueta, A., Aznar, M.A., Quetglas, J.I., et al. (2016). Cancer Immunotherapy with Immunomodulatory Anti-CD137 and Anti-PD-1 Monoclonal Antibodies Requires BATF3-Dependent Dendritic Cells. Cancer Discov 6, 71–79.

Sanjuan, M.A., Milasta, S., and Green, D.R. (2009). Toll-like receptor signaling in the lysosomal pathways. Immunol Rev 227, 203–220.

Saveanu, L., Carroll, O., Weimershaus, M., Guermonprez, P., Firat, E., Lindo, V., Greer, F., Davoust, J., Kratzer, R., Keller, S.R., et al. (2009). IRAP identifies an endosomal compartment required for MHC class I cross-presentation. Science 325, 213–217.

Savina, A., and Amigorena, S. (2007). Phagocytosis and antigen presentation in dendritic cells. Immunol Rev 219, 143–156.

Saxena, M., and Bhardwaj, N. (2018). Re-Emergence of Dendritic Cell Vaccines for Cancer Treatment. Trends Cancer 4, 119–137.

Schindelin, J., Arganda-Carreras, I., Frise, E., Kaynig, V., Longair, M., Pietzsch, T., Preibisch, S., Rueden, C., Saalfeld, S., Schmid, B., et al. (2012). Fiji: an open-source platform for biological-image analysis. Nat Methods 9, 676–682.

Serreze, D.V., Chapman, H.D., Varnum, D.S., Gerling, I., Leiter, E.H., and Shultz, L.D. (1997). Initiation of autoimmune diabetes in NOD/Lt mice is MHC class I-dependent. J Immunol 158, 3978–3986.

Shen, L., Sigal, L.J., Boes, M., and Rock, K.L. (2004). Important role of cathepsin S in generating peptides for TAP-independent MHC class I crosspresentation in vivo. Immunity 21, 155–165.

Shvets, E., and Elazar, Z. (2009). Flow cytometric analysis of autophagy in living mammalian cells. Methods Enzymol 452, 131–141.

Snyder, A.G., Hubbard, N.W., Messmer, M.N., Kofman, S.B., Hagan, C.E., Orozco, S.L., Chiang, K., Daniels, B.P., Baker, D., and Oberst, A. (2019). Intratumoral activation of the necroptotic pathway components RIPK1 and RIPK3 potentiates antitumor immunity. Sci Immunol 4.

Stober, D., Trobonjaca, Z., Reimann, J., and Schirmbeck, R. (2002). Dendritic cells pulsed with exogenous hepatitis B surface antigen particles efficiently present epitopes to MHC class I-restricted cytotoxic T cells. Eur J Immunol 32, 1099–1108.

Turner, M.D., Nedjai, B., Hurst, T., and Pennington, D.J. (2014). Cytokines and chemokines: At the crossroads of cell signalling and inflammatory disease. Biochim Biophys Acta 1843, 2563–2582.

Wolfers, J., Lozier, A., Raposo, G., Regnault, A., Thery, C., Masurier, C., Flament, C., Pouzieux, S., Faure, F., Tursz, T., et al. (2001). Tumor-derived exosomes are a source of shared tumor rejection antigens for CTL cross-priming. Nat Med 7, 297–303.

Wu, S.J., Niknafs, Y.S., Kim, S.H., Oravecz-Wilson, K., Zajac, C., Toubai, T., Sun, Y., Prasad, J., Peltier, D., Fujiwara, H., et al. (2017). A Critical Analysis of the Role of SNARE Protein SEC22B in Antigen Cross-Presentation. Cell Rep 19, 2645–2656.

Yatim, N., Jusforgues-Saklani, H., Orozco, S., Schulz, O., Barreira da Silva, R., Reis e Sousa, C., Green, D.R., Oberst, A., and Albert, M.L. (2015). RIPK1 and NF-kappaB signaling in dying cells determines cross-priming of CD8(+) T cells. Science 350, 328–334.

Zhang, N., and Bevan, M.J. (2011). CD8(+) T cells: foot soldiers of the immune system. Immunity 35, 161–168.

## References

Komatsu, M., et al., Impairment of starvation-induced and constitutive autophagy in Atg7-deficient mice. J Cell Biol, 2005. 169(3): p. 425–34.

Kundu, M., et al., Ulk1 plays a critical role in the autophagic clearance of mitochondria and ribosomes during reticulocyte maturation. Blood, 2008

Martinez, J., et al., Molecular characterization of LC3-associated phagocytosis reveals distinct roles for Rubicon, NOX2 and autophag proteins. Nat

Martinez, J., et al., Noncanonical autophagy inhibits the autoinflammatory, lupus-like response to dying cells. Nature, 2016. 533(7601): p. 115–9.

